# The divergence of DHN-derived melanin pathways in *Metarhizium robertsii*

**DOI:** 10.1101/2024.07.25.605210

**Authors:** Linan Xie, Yang Liu, Yujie Zhang, Kang Chen, Qun Yue, Chen Wang, Baoqing Dun, Yuquan Xu, Liwen Zhang

## Abstract

The important role of dihydroxynaphthalene-(DHN) melanin in enhancing fungal stress resistance and its importance in fungal development and pathogenicity are well-established. This melanin also aids biocontrol fungi in surviving in the environment and effectively infecting insects. However, the biosynthetic origin of melanin in the biocontrol agents, *Metarhizium* spp., has remained elusive due to the complexity resulting from the divergence of two DHN-like biosynthetic pathways. Through the heterologous expression of biosynthetic enzymes from these two pathways in baker’s yeast *Saccharomyces cerevisiae*, we have confirmed the presence of DHN biosynthesis in *M. roberstii*, and discovered a novel naphthopyrone intermediate that can produce a different type of pigment. These two pigment biosynthetic pathways differ in terms of polyketide intermediate structures and subsequent modification steps. Stress resistance studies using recombinant yeast cells have demonstrated that both DHN and its intermediates confer resistance against UV light prior to polymerization; a similar result was observed for its naphthopyrone counterpart. This study contributes to the understanding of the intricate and diverse biosynthetic mechanisms of fungal melanin and has the potential to enhance the application efficiency of biocontrol fungi such as *Metarhizium* spp. in agriculture.

## 1. Introduction

Melanins are ubiquitous pigments found in all kingdoms of life, mostly involved in protection of abiotic and biotic stress such as UV light or ionizing radiation, toxic substances, and oxidative stress.^1-3^ 1,8-Dihydroxynaphthalene (DHN)-melanin is one dominant type in fungi especially ascomycetes and hemimycetes, and is frequently involved in stress resistance and pathogenicity.^4-7^ In both saprophytes and pathogens, the accumulation of DHN melanin in reproductive structures such as conidia can bolster cell-wall strength and protect the genetic materials from damage; lacking the DHN melanin can lead to the failure of spore development.^8-9^ In pathogenic fungi, the DHN melanin is essential to withstand the high turgor pressure in appressoria during infection.^3, 10^ Blocking its synthesis pathway can significantly reduce or even eliminate the infection ability to plants and animals. Therefore, the biosynthesis mechanism of DHN melanin is one key to understand the resistance of pathogenic fungi to stress and their interaction with host.

DHN melanin is a kind of brown or black macromolecular polymer formed by oxidative polymerization of phenolic monomers. It is insoluble and resistant to degradation by acids, which is why their structures are difficult to analyze and are not fully understood.^11^ The first key intermediate product of this pathway is 1,3,6,8-tetrahydroxynaphthalene (T4HN), which is then sequentially converted to scytalone, 1,3,8-tetrahydroxynaphthalene (T3HN), vermelone and DHN by 1,3,4-trihydroxynaphthalene reductase (THNR) and scytalone dehydratase (SCD).^12-14^DHN as the last monomer is polymerized into melanin by phenol oxidase, peroxidase, laccase or catalase.^15^ DHN biosynthetic pathway widely exists in fungi and is very conservative, indicating its significance. Biosynthesis of T4HN, the indispensable precursor to DHN melanin, is similar but divergent in fungi. The initiation of T4HN synthesis in fungi is catalyzed by non-reducing polyketide synthase (nrPKS), referred to as T4HN synthase in this paper. At present, three T4HN pathways have been identified: hepta-, hexa- and penta-ketone pathway. The representative heptaketide, YWA1 (2-malonyl-T4HN), can be synthesized by the nrPKS (Alb1) from *Aspergillus fumigatus*,^16^ while WdPKS1 from *Wangiella dematitidis* produces hexa-ketone 2-acetyl-T4HN (AT4HN).^17^ These two intermediates require serine hydrolase Ayg1p or WdYg1p, respectively, to deacylate the side-chain and generate T4HN.^18-19^ ClPKS1 from *Colletotrichum orbiculale* can directly synthesize penta-ketide T4HN.^20-21^ Domain swapping demonstrated the bifunctional thioesterase domain of ClPKS1 catalyzing both cyclization and deacetylation of the enzyme-bound hexaketide substrate to produce the penta-ketide product.^22^ The PfmaE identified in the plant endophytic fungus *Trichoderma ficus* was the second example that can directly synthesize T4HN.^9^

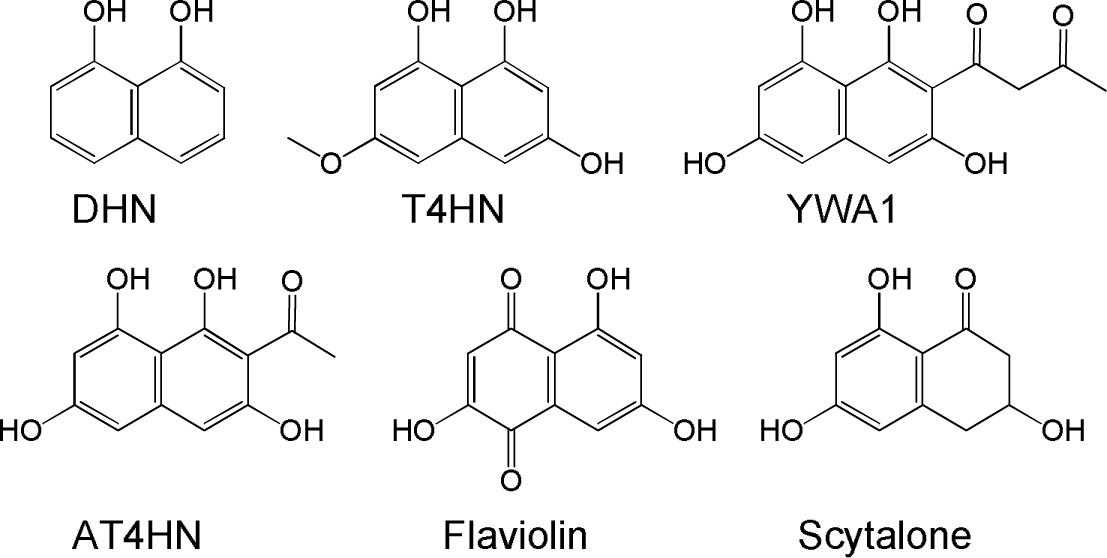

Entomopathogenic fungi have been used for biocontrol of insect pests for many decades. However, the efficacy of such fungi in field trials is often inconsistent, mainly due to environmental stresses, such as UV radiation, temperature extremes, and desiccation.^15^ The factors related to fungal virulence and environmental stress responses have close connections with fungal biocontrol efficacy.^23-24^ *Metarhizium* species are among the most abundant entomopathogenic fungi. Species such as *M. roberstii* and *M. anisopliae* are the most widely used biocontrol fungus in agricultural and forestry pest control, and are considered as model organisms to study the host-pathogen interaction mechanism.^25-27^ Most *Metarhizium* fungi produce light or dark green pigments, which were shown to protect the cell from abiotic stresses (e.g., ultraviolet radiation and heat shock), and affect pathogenicity.^28-29^ Transferring the DHN melanin pathway genes (PKS, SCD, and THNR genes from *Alternaria alternata*) into *M. anisopliae* significantly enhanced the stress resistance and pathogenicity.^15^

However, the endogenous melanin pathway of *Metarhizium* species is still a mystery. On one hand, using DHN melanin synthesis inhibitor tricyclazole failed to cause phenotypic change in the *M. anisopliae* strain ARSEF 2575 (now classified as *M. robertsii*), indicating absence of DHN pathway and presence of a unique, yet undetermined, melanin synthesis pathway.^24, 29^ On the other hand, in the sequenced genomes of seven *Metarhizium* species except for the albino *M. album*, two nrPKSs (*Pks1* and *Pks2*) were found to be highly homologous to the T4HN synthase genes.^24, 30^ These two genes are evolved from replication and differentiation events, but their expression pattern and biological functions are different.^28^ The *Pks2* clusters are highly conserved with the DHN-melanin pathway, whereas the disruption of these PKS genes did not change the color; no related product was identified when the *MrPks2* gene from *M. roberstii* was heterologously expressed in *Aspergillus nidulans*.^28^ Thus, the *Pks2* genes seemed not to be involved in melanin synthesis. The *Pks2* genes were activated on insect cuticle, and their disruption mutants showed prolonged formation of appressoria and reduced insecticidal ability.^24, 28^ In contrast, *Pks1* genes are highly expressed during conidiation but not on insect cuticle, and their disruption led to the brownish red color instead of dark green, indicating the key role in melanin/pigment synthesis. Heterologous expression of *MrPks1* gene produced the anthraquinone compound 1-acetyl-2,4,6,8-tetrahydroxy-9,10-anthraquinone.^28^ However, no report showed that this compound is related to pigment synthesis.

Most of the previous studies on DHN melanin used gene disruption, deletion or overexpression in the original fungal strain, especially for biological function studies. As an important complement to these methods, heterologous expression has the advantage of circumventing the restriction of the native producer strain, and moreover, the impact of microbial culture conditions, the unknown outcomes of gene cluster activation, complex product isolation and identification. A few studies used heterologous host such as *A. nidulans* or *A. oryzae* to elucidate the biosynthesis pathway.^20, 28, 31,32^ However, these hosts have complex secondary metabolism including their own pigment biosynthesis pathways, which may interfere the heterologous pathways. Therefore, we tried to investigate the divergent biosynthesis pathways of PKS1 and PKS2 from *M. robertsii* through heterologous reconstruction of the natural and combinatorial biosynthetic pathways in yeast (*Saccharomyces cerevisiae* BJ5464-NpgA), a host does not produce melanin and has relative “clean” secondary metabolic background. This can help us to approach the authentic structure and function of the DHN melanin and its intermediates. The result is not only helpful to understand the mechanism of fungi adapting to the environment and their hosts, but also of great value for the development of efficient fungal pesticides.

## 2. Materials and Methods

### Chemicals

All substrate, except compound **1-3**, were purchased from MilliporeSigma (St. Louis, MO, United States). Compound **1-3** were isolated from the recombinant *S. cerevisiae* BJ5464-NpgA expressing the fungal polyketide synthases MrPks1.

### Strains, plasmids, and culture conditions

The fungus *M. robertsii* ARSEF2575 was kindly provided by Prof. Chengshu Wang from Center for Excellence in Molecular Plant Sciences, Shanghai Institute of Plant Physiology and Ecology, CAS. The strain was cultured on potato dextrose agar (PDA) plates at 28 °C, and the collected mycelia were used for genomic DNA extraction. Spores were collected after 10 days of growth by flooding the plates with sterile 0.1% Triton-X100, and allowed to grow in potato-dextrose broth (PDB) (Difco, Sparks, MD, USA) medium before the RNA isolation at 28 °C.

*Escherichia coli* DH5α and plasmid pJET1.2 (Thermo Fisher) were used for routine cloning and sequencing. *Saccharomyces cerevisiae* BJ5464-NpgA (*MAT*α *ura3-52 his3-*Δ*200 leu2-*Δ*1 trp1 pep4::HIS3 prb1* Δ*1.6R can1 GAL*) was used as the host for expression vectors based on plasmids YEpADH2p-URA, YEpADH2p-TRP and YEpADH2p-LEU. The construction of plasmids and the cultivation of recombinant *S. cerevisiae* BJ5464-NpgA strains were carried out as previously described^33-37^ in three independent *S. cerevisiae* transformants for each recombinant yeast strain, and fermentations with representative isolates were repeated at least two times. For biotransformation assays, substrates (10 mg/L, final concentration) in methanol were supplemented to the culture when it reached an OD600 of 0.6, and incubation was continued at 30 °C with shaking at 220 rpm for an additional two days. After that, the fermentation broth was filtered, and the filtrate was neutralized to pH 7.0 before the extraction using an equal volume of ethyl acetate. *Aspergillus nidulans* A1145 (*pyrG89;pyroA4;nkuA::argB;riboB2*) was also used as the host for gene expression as described in previous studies.^9,28^

### RNA isolation and RT-PCR

The mycelia of *M. robertsii* cultured on SDY (Sabouraud dextrose medium with yeast extract) or PDA (potato dextrose agar) plates for 6 days were collected. Total RNAs from the mycelia were extracted using the Trizol (Invitrogen, Carlsbad, CA, United States). The RNAs were reversely transcribed into First strand cDNA with Superscript II reverse transcriptase (TaKaRa Bio. Inc., Kusatsu, Japan) following the manufacturer’s protocol. Primers were designed from the start and stop codon of the genes (Table S5). PCR amplification was performed on a BioRad Thermocycler using Phanta®Max Super-Fidelity DNA Polymerase (Vazyme Biotech Co., Ltd, Nanjing, China). Vectors were constructed using the CloneExpress Kit (Vazyme Biotech Co., Ltd, Nanjing, China), based on the described vector system. The designated genes were inserted between the NdeI-PmeI sites of the YEpADH2p expression vectors featuring different selectable markers. All newly constructed plasmids were verified by DNA sequencing.

### Chemical characterization

The detection, isolation and chemical characterization of products were followed previously reported procedures.^33-37^ Briefly, extracts were analyzed by liquid chromatography-mass spectrometry (LC-MS), and products were isolated from scaled-up fermentations (1-10 L, depending on yield). LC-MS runs were performed on an Agilent 1290 Infinity II HPLC (Santa Clara, CA, United States) coupled with an Agilent InfinityLab single quadrupole mass selective detector operating in negative ion mode. The HPLC was equipped with an Agilent Eclipse Plus C18 RRHD column (50 mm x 2.1 mm id, 1.8 µm particle size), and eluted with a linear gradient of 10-50% of acetonitrile-water over 4 min, 50-95% for 4 min, 95% acetonitrile-water for 2 min and drop down to 10% in 1 min at a flow rate of 0.35 mL/min. Semi-preparative HPLC was performed on an Agilent Eclipse XDB-C18 reversed-phase column (5 µm, 9.4 mm × 250 mm) using an Agilent 1260 Infinity II instrument at the flow rate of 2 mL/min. HRESIMS spectra were recorded on an Agilent QTOF 6530 instrument operated in negative ion mode using capillary and cone voltages of 3.6 kV and 40-150 V, respectively. The collision energy was optimized from 15 to 50 V. For accurate mass measurements the instrument was calibrated each time using a standard calibration mix (Agilent) in the range of *m*/*z* 100-1900. ^1^H, ^13^C, HSQC, and HMBC NMR spectra were recorded on a Bruker Avance III 400 MHz spectrometer. Chemical shift values (*δ*) are given in parts per million (ppm), and the coupling constants (*J* values) are in Hz. Chemical shifts were referenced to the residual solvent peaks of DMSO-*d*_6_.

### Tolerance to stress

Tolerance to heat shock was investigated by incubating yeast expressing designated genes at 45, 50 or 55 LJ for 2 h then transferred to 30 LJ to continue growth. The growth was checked every 2 h. The tolerance to UV-B radiation was determined by measuring the growth of yeast colonies following exposure to 400 μW cm^-2^ of Quaite-weighted UV-B radiance. Exposures of 10 and 30 min afforded total doses of 0.24 and 0.72 J cm^-2^, respectively. After irradiation, yeast (OD 0.8) were serial diluted at 10-fold on SC minimal dropout (-Trp, -Ura, -Leu) agar plates (Clontech) at 30 LJ.

## 3. Results

### 3.1 MrPKS2 synthesized T4HN in S. cerevisiae

To identify the products of MrPKS1 and MrPKS2, the intron-free *MrPks1* and *MrPks2* open reading frames were expressed in *S. cerevisiae* BJ5464-NpgA, a host adopted to reconstitute fungal polyketide biosynthetic pathways.^33-34, 38-40^ In previous study, heterologous expression of *MrPks2* in *A. nidulans* showed no product peak.^28^ In contrast, the yeast strain successfully produced the major product of MrPKS2 (Fig. 1A). The mass-to-charge ratio (*m*/*z*) ([M-H]^-^ ions) of this product was 205.0142 atomic mass unit (amu), consistent to a molecular formula of C_10_H_6_O_5_ (calcd. *m*/*z* 205.0136, mass error 2.4 ppm). This formula and the HRMSMS spectrum were in accordance with flaviolin that can be derived from spontaneous oxidization of T4HN (Fig. 1B). Similarly, treatment of *C. lagenarium* with the DHN pathway inhibitor, tricyclazole, resulted in the accumulation of flaviolin instead of T4HN.^22^ Thus, our result shows that MrPKS2 is a T4HN synthase likely belonging to the penta-ketide class with ClPKS1 and PfmaE.^9, 20-21^

**Figure 1.**
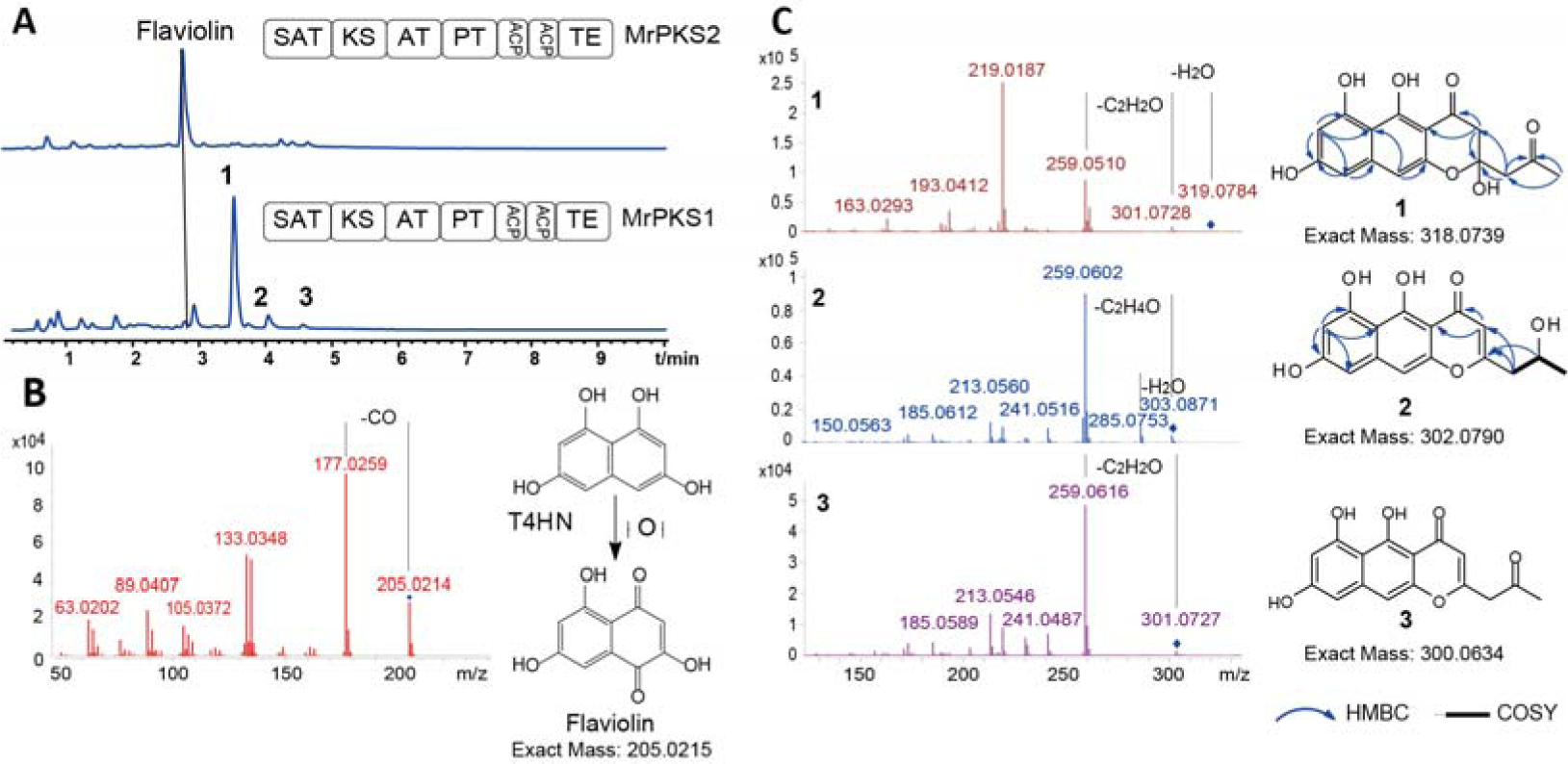
The divergent products produced by MrPKS1 and MrPKS2. A. Product profiles (reversed phase HPLC-UV traces recorded at 280 nm) of *S. cerevisiae* BJ5464-NpgA expressing the indicated PKSs from *M. robertsii*; the domain architectures of both PKSs were illustrated with the round rectangles representing designated domains. The abbreviation of domains were SAT: starter acyltransferase, KS: ketoacyl synthase, AT: the acyltransferase, PT: product template, ACP: acyl carrier protein domain and TE: thioesterase. B. Structures of T4HN and flaviolin and HRESIMS/MS spectrum of flaviolin. C. The HRESIMS/MS spectra and the structures of **1-3**.

### 3.2 MrPKS1 synthesized new octaketides in S. cerevisiae

Different with *MrPks2*, the recombinant yeast strain expressing *MrPks1* afforded three product peaks, **1-3** (Fig. 1A). The positive mode HRESIMS spectra of the **1-3** displayed [M+H]^+^ ions at *m*/*z* 319.0784, 303.0871 and 301.0727, corresponding to the molecular formula of C_16_H_14_O_7_ (mass error -10.30 ppm), C_16_H_14_O_6_ (mass error 0.66 ppm) and C_16_H_12_O_6_ (mass error 4.98 ppm), respectively (Fig. 1C). As the major product of MrPKS1, **1** showed *m*/*z* value in agreement with the mass of an octaketide with new structure. Its structure was elucidated by NMR (Fig. S1 and Table S1). The ^1^H NMR data for compound **1** (Table S1-1) revealed the presence of three aromatic proton signals [δ_H_ 6.19 (1H, s), 6.39 (1H, s), 6.39 (1H, s)] and one singlet methyl signal (δ_H_ 2.17, s). The ^13^C NMR spectrum (SI Table S1-2), with the aid of HSQC, showed 16 resonances ascribed to a methyl, two methylenes, three methines, and ten quaternary carbons. Analysis of 1D and 2D NMR data (Fig. S1 and Table S1) showed high similarities between **1** and YWA1,^41^ except for the extra acetyl group. The position of the acetyl group was determined at C-11 according to the HMBC correlations from H_2_-11 and H_3_-13 to the ketone carbonyl at δ_C_ 205.0. All of the NMR signals and correlations are consistent with the structure of **1** (Fig. 1C), which is equivalent to attaching one more keto-unit to the structure of YWA1, the heptaketide in DHN melanin pathway. YWA1 can be reversibly convertible to 2-malonyl-T4HN. Similarly, **1** may also be converted from a similar intermediate (Fig. 1C).

MrPKS1 is the first reported natural T4HN synthase producing octaketide. MrPKS1 maintained precise control of cyclization regioselectivity. Octaketides that contain alternative cyclization patterns was not detected. Similar compounds except for YWA1 have also been reported, i.e., the intermediate (**5**) in the biosynthesis of pre-bikaverin by PKS4 from *Giberella fujikuroi* (Fig. 2C).^42-43^ The compound **5** has one more keto-unit attached to the end of **1**. It is highly possible that the poly-β-keto processing pathway of **1** is identical to those during YWA1 and pre-bikaverin synthesis catalyzed by *A. nidulans* wA synthase and *G. fujikuroi* PKS4 synthase, respectively. In brief, cyclization of the first benzene ring is facilitated by the aldol condensation reaction between C-2 and C-7, which requires synthesis of the full length polyketide chain. This is followed by Claisen-like condensation between C-10 and C-1 to generate the bicyclic T4HN scaffold (Fig. 2C).^42-43^ Attack of C-13 carbonyl by the nucleophilic C-9 phenol forms the tricyclic naphthopyrone.

**Figure 2.**
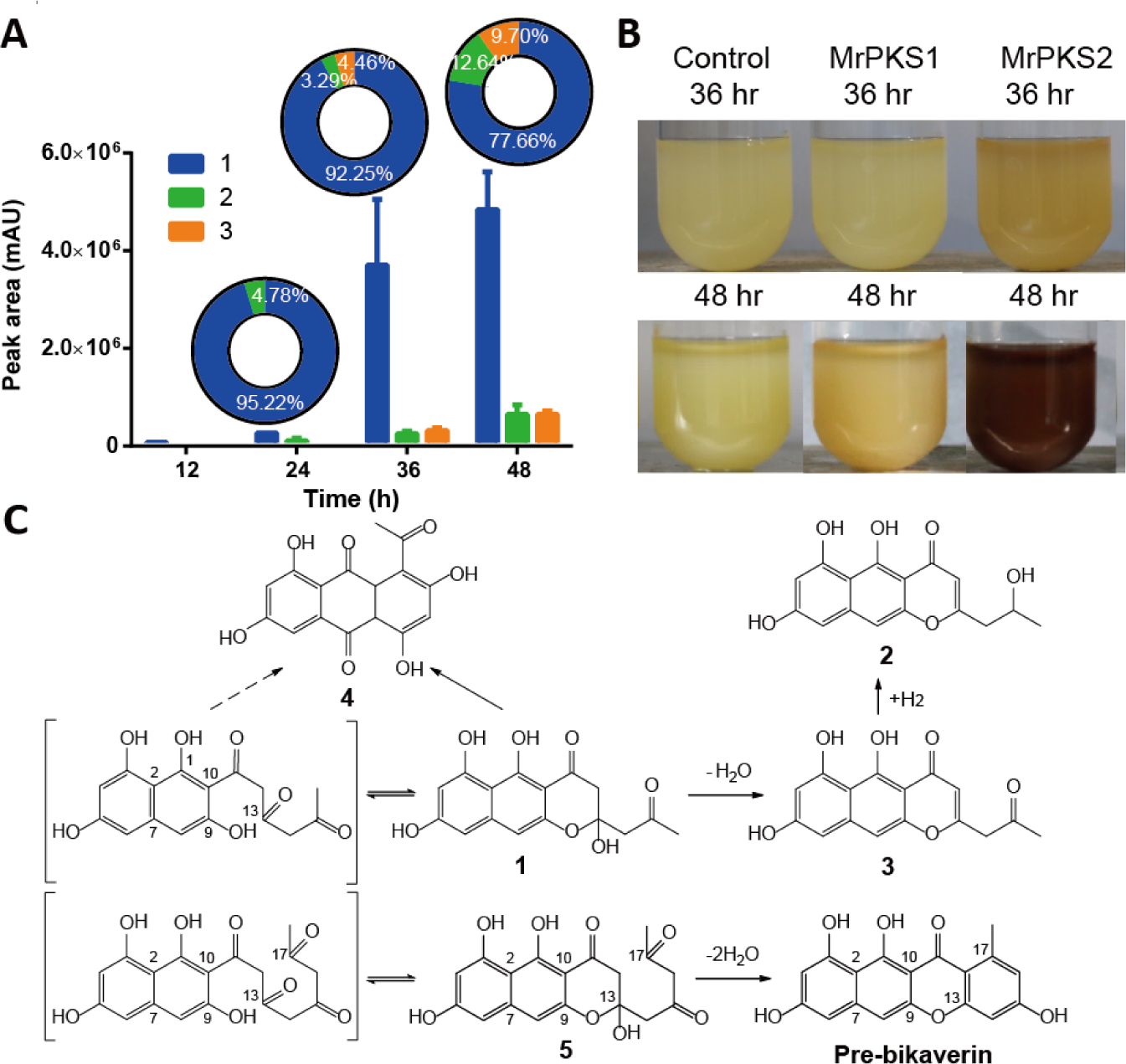
Time course of MrPKS1 product profiles recorded at 12, 24, 36 and 48 hrs. A. Peak area (bar graph) and relative percentage (ring graph) of compound **1-3** calculated from the reversed phase HPLC-UV traces recorded at 280 nm. B. Photo of *S. cerevisiae* expressing the indicated PKSs from *M. robertsii* after 36 or 48 hrs. The control strain expressed the plasmid without PKS genes. C. The proposed poly-β-keto processing pathway of **1-4**.

The highly similar high resolution tandem mass (HRMS/MS) fragmentation pattern of **1**-**3** demonstrated their structure similarity (Fig. 2B and Table S2). Similar to **1**, the structure of **2** can be elucidated by the NMR spectra and the HRMS/MS spectrum (Fig. 2B, Fig. S1 and Table S1). In addition to the same aromatic ring skeleton with compound **1**, the presence of 2-(2-hydroxypropyl)-γ-pyrone group can be confirmed in compound **2** according to the HMBC correlations from H_2_-11 to C-2 and C-3, from H-3 to C-4 and C-4a, associated with ^1^H-^1^H COSY correlations from H-12 to H_3_-13 and H_2_-11 (more details in supporting information). Therefore, the structure of compound **2** was determined as shown in Fig. 1.

According to the structures of **1** and **2**, the structure of **3** can be proposed. The hydroxyl group at C13 position of **1** is easily dehydrated to form **3**, which can be demonstrated by the presence of 301.0728 fragment ion in the high resolution tandem MS (HRMS/MS) spectra of **1** (mass error -5.31 ppm) (Fig. 2B). This ion was 18 amu less than the parent ion of **1**, and was in accordance to the *m*/*z* of **3** (301.0727, [M-H]^+^). The carbonyl group of **3** at C15 position was reduced to hydroxyl group to form **2**. This was corresponding to the near identical fragmentation pattern in HRMS/MS spectra of **2** and **3**, except that the **2**’s spectra had the –18 amu fragment ion (*m*/*z* 285.0753, the loss of hydroxyl group at C15).

### 3.3 The reactivity of MrPKS1 products

The time course study showed that **1** can be detected after 24 h fermentation with the *MrPks1* expressed, reached to a plateau at around 36 h and remained a similar level until 48 h (Fig. 2A). Photos taken at the same periods were in accordance with the product profiles, which showed color difference at 36 hrs and 48 hrs with yellow to brownish yellow color (Fig. 2B). The time course profiles also showed the emergence of minor products, **2** and **3**. The dehydration from **1** to **3** and the reduction from **3** to **2** were likely to occur spontaneously. The crude extract of MrPKS1-expressing yeast showed increased amount of **2** and **3** after 1 week (Fig. S2A-B); the purified **1** product in DMSO also gave rise to **2** and **3** (Fig. S2C-D).

Expression of *MrPks1* in *A. nidulans* A1145 resulted in the product **4** in previous study^28^ as well as in ours (Fig. 2C and Fig. S3). Trace amount of **4** was also detected in the yeast crude extract expressing *MrPks1* with the same retention time and HRMS/MS spectrum as those detected in *A. nidulans*. The control strains of *A. nidulans* and yeast both lack this product peak, which confirmed the origin of **4** to be MrPKS1. We also conducted the time course study of MrPKS1 expressed in *A. nidulans* (Fig. S3), which showed the occurrence of both **1** and **4** after 3-d fermentation. However, the **1** disappeared after 5 d in *A. nidulans* but still accumulated in yeast as in Fig. 1. In a word, these results showed that **1** is closer to the authentic product of MrPKS1. Although the detailed routes from **1** to the other forms are still unknown, the high reactivity of MrPKS1’s products can be demonstrated.

### 3.4 Synteni analysis of loci encoding the T4HN synthases

Fungal genomes prefer to encode the genes that collaborate to synthesize a secondary metabolite in adjacent loci, so called biosynthetic gene cluster (BGC). Thus the gene composition of BGC provides insights into the biosynthesis pathway, and the gene rearrangement gives clues to evolution process. To better understand the divergence of *Pks1* and *Pks2* pathways, we performed phylogenetic and syntenic analysis of the homolog BGCs in ascomycete fungi that has been reported to biosynthesize DHN melanin. 19 BGCs containing homologous T4HN synthases were submitted to CORASON.^44^ Sequences from 15 species were aligned and used to construct the phylogenetic tree with bootstrap values > 88% except for PfmaE (Fig. 3). Since the BGCs of 6 *Metarhizium* species are highly conserved, three representative species, the broad insecticidal *M. robertsii* and *M. anisopliae* as well as the host-specific *M. acridum*, were selected in this analysis. In addition to the T4HN synthases, homologs of tailoring enzymes were also searched with e-value cut-off of 10^-1^. In accordance to the previous study using ketosynthase domain sequences to construct the tree,^28^ the PKS1s and PKS2s from the *Metarhizium* species formed a clade with the taxonomically close Hypocreales fungus, *Fusarium gramineanum*, with high confidence (96% bootstrap value) (Fig. 3). This clade was divided into two well-supported sub-clades corresponding to PKS1 and PKS2, respectively (Fig. 3). The T4HN synthase from *F. gramineanum* is more similar to PKS1, suggesting a closer relationship of PKS1 to the common ancestor PKS. Basal to this clade was the clade of the reference BGCs of Alb1 and wA from the Eurotiomycetidae *A. fumigatus* and *A. nidulans*. The Alb1 BGC is composed of seven genes encoding proteins involved in DHN-melanin biosynthesis: from downstream to upstream were the T4HN synthase (Alb1), SCD (Arp1), THNR (Arp2) and serine hydrolase (Ayg1) that required for DHN synthesis, as well as two oxidases/laccases (Abr1 and Abr2) and a transcriptional factor.^16^ Comparison of the genomic context of PKS1 and PKS2 to that of Alb1 indicates that they separately “inherited” different tailoring enzymes after the duplication event: PKS2 took THNR and SCD while PKS1 adopted the laccase/oxidase (MLac1) that is homologous to Abr2. The serine hydrolase was no longer needed since PKS2 can directly synthesize T4HN. The “remnant” of *Abr1* gene homolog in the BGC from *M. anisopliae* suggests the gradual loss of this gene. Further examination of the other remote clades/BGCs showed that *Abr2* (*MLac1*) and *THNR* were two genes that frequently occurred, which also indicates their common origin and departure in *Metarhizium* species. Deletion of the *abr2* gene in *A. fumigatus* changed the gray-green conidial pigment to a brown color compared with wild-type conidia,^45^ suggesting that Abr2 may be the laccase that polymerize DHN. Thus it is reasonable that the PKS2 BGC “cross-talk” with the PKS1 BGC and use the MLac1 to conduct the polymerization.

**Figure 3.**
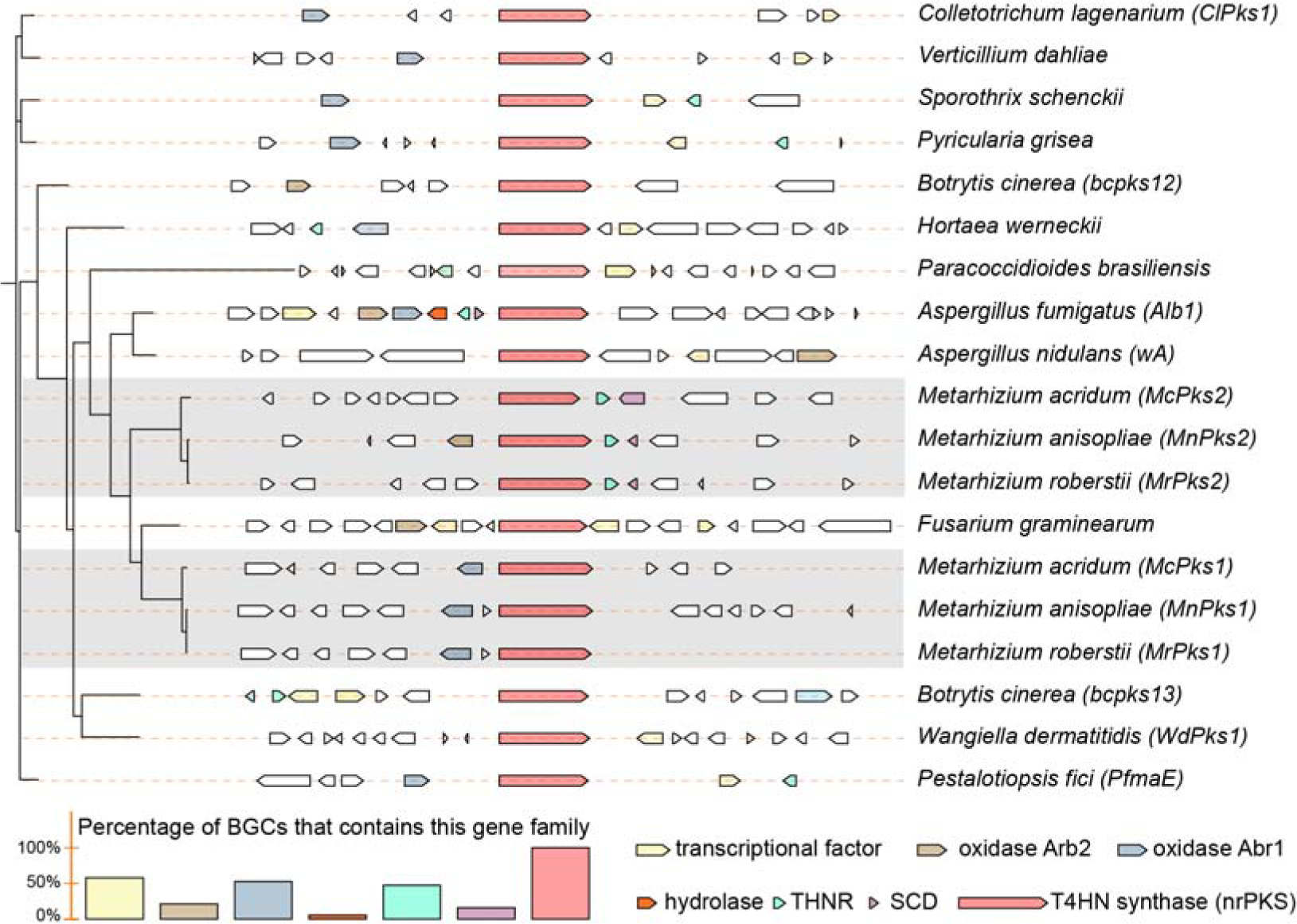
Phylogenetic and syntenic analysis of the PKS1, PKS2 from *M. robertsii*, *M. anisopliae* and *M. acridum*, as well as other T4HN synthases from ascomycete fungi that has been reported to biosynthesize DHN or DHN-like melanins. The phylogenetic tree (left) was constructed based on the amino acid sequences of the T4HN synthases. The Genbank accession number for each full-length amino acid sequence of each PKS and the references were included in Table S3. The clade containing the *Metarhizium* PKS1 and PKS2 were shaded. The scale bar corresponds to the estimated number of amino acid substitutions per site. Schematic maps (middle) of the gene clusters containing PKS1, PKS2, or their homologs were constructed using CORASON. Colors in the inset legend represent genes in the cluster; white indicate genes that are not homologous to each other (cut-off e value was 10^-1^). The fungal species are indicated right of the gene cluster maps. Percentage values of the BGCs that contain specific gene family were shown in the lower left bar graph.

In addition to the proteins present in the Alb1 BGC, the Pks1 BGC also includes a gene encoding a protein with an EthD domain (Pfam07110); this domain is involved in the degradation of ethyl tert-butyl ether.^46^ Previous study has shown that in the genomic context of MrPKS1, only MrPKS1, MrEthD and MLac1 appear to involve in the dark greenish color of *M. robertsii*: the mutants with MrPKS1 disrupted produced light red conidia; while the mutants with MrEthD or MLac1 disrupted had dark red conidia.^46^ It is very likely that MLac1 can polymerize the phenolic product of PKS1, while EthD participate/aid this process. However, whether these three genes are sufficient to the biosynthesis of melanin/pigment (i.e., as a DHN bypass similar to the wA BGC), or in another case the **1** intermediate convergently resulted in DHN still remains a question.

### 3.5 The biosynthesis of DHN by the MrPks2 gene cluster

Based on the synteni analysis, we can hypothesize that the *MrPks2* gene cluster is capable to biosynthesize DHN, which could be further polymerized into melanin through the “cross-talk” with the MLac1 encoded in the *MrPks1* gene cluster. To test this hypothesis, the intron-free *MrTHNR* and *MrSCD* open reading frames were both cloned from the reverse-transcribed cDNA; this cDNA was prepared during cuticle infection.^24^ Sequencing of the cloned fragments confirmed the presence of 2 introns in both *MrTHNR* and *MrSCD*, resulted in 109 bp and 150 bp length shorter compared to the genes encoded in genomic DNA (Fig. S4). Then the *MrTHNR* and *MrSCD* were co-expressed with *MrPks2* separately or collectively in *S. cerevisiae* BJ5464-NpgA. Strains expressing *MrPks2* and *MrTHNR* yielded only trace amount of flaviolin; instead, produced a major product with *m*/*z* ([M-H]^-^ ion) of 193.0494 that is correspondent to scytalone (C_10_H_10_O_4_, calcd. *m*/*z* 193.0506, mass error -6.22 ppm) (Fig. 4). The structure of this product was further elucidated by analyzing the NMR spectroscopic data of the purified compounds (Fig. S5, Table S4). The expected product, DHN, was also detected in yeast expressing *MrPks2*, *MrTHNR* and *MrSCD*, after 36 h and 48 h fermentation (Fig. 4). The HPLC-HRMS graphs clearly showed the presence of a product peak with [M-H]^-^ of 159.0430 (molecular formula C_10_H_8_O_2_, calcd. *m*/*z* 159.0446, mass error 2.4 ppm). The HRMS/MS spectra (Fig. 4C) and the identical retention time in HPLC graph (Fig. S5) showed that this product is highly possible to be DHN. Therefore, the BGC of MrPKS2 is fully functional to generate DHN and participates in the melanin biosynthesis; the reason why expressing *MrPks2* in *A. nidulans* did not show detectable product^28^ may be because the T4HN produced by MrPKS2 was utilized by the host melanin pathways.

**Figure 4.**
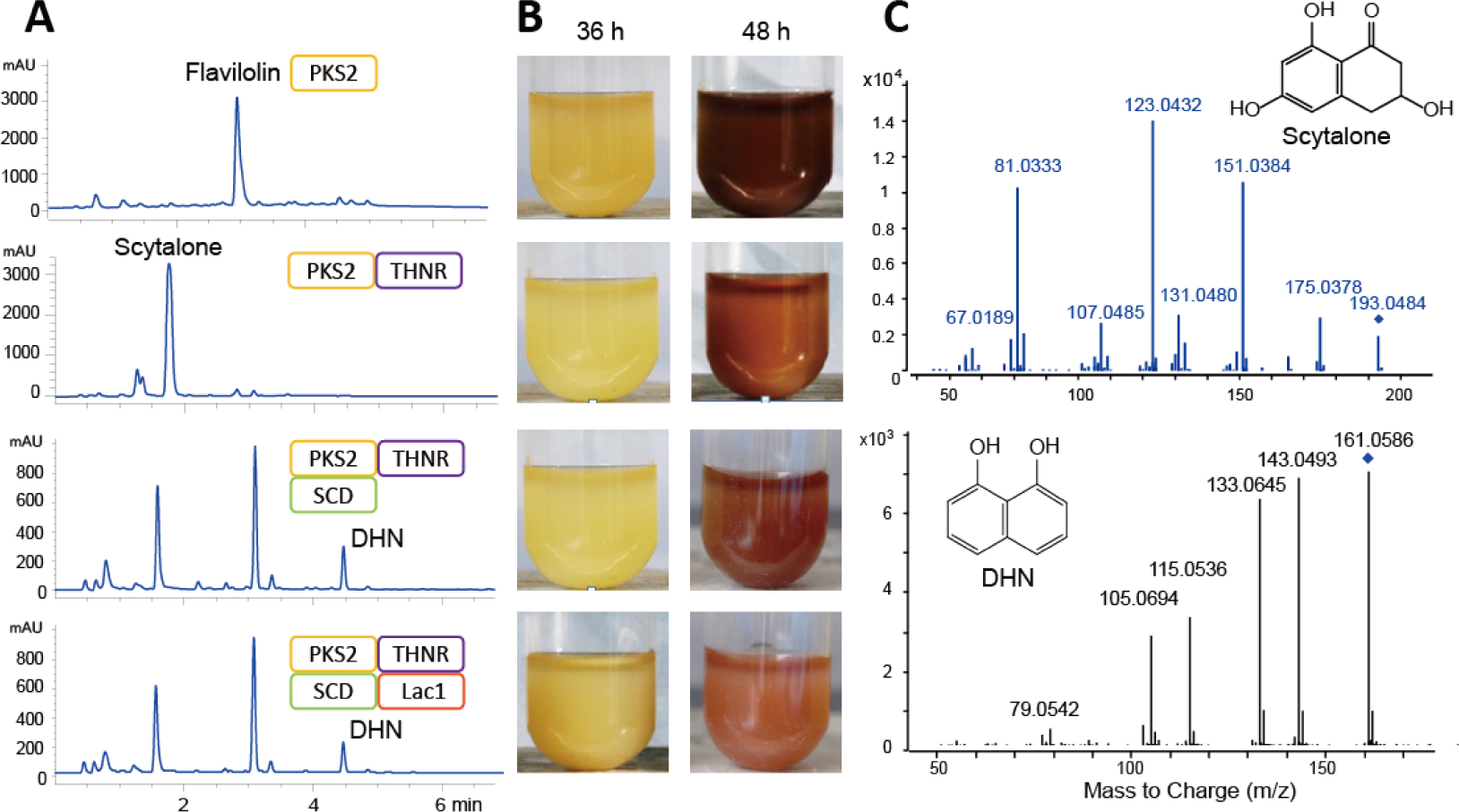
Recombination of *Metarhizium* DHN melanin biosynthesis pathway in yeast. A. Product profiles (reversed phase HPLC-UV traces recorded at 280 nm) of *S. cerevisiae* BJ5464-NpgA expressing the indicated genes from *M. robertsii*. B. The colors of yeast fermentations at 36 or 48 hr. C. HRESIMS/MS spectra of scytalone and DHN; the structures were illustrated.

The reason that the disruption of *MrPks2* gene did not cause color change to *Metarhizium* species is due to its low expression during saprophytic growth. According to RT-PCR (reverse transcription polymerase chain reaction) result (Fig. S4), the genes *MrTHNR* and *MrSCD* were strictly regulated and were only transcribed during infection. This is also the reason why blocking the DHN biosynthesis did not cause physiological change to the *Metarhizium* spp. However, the *MrPks2* gene showed transcription on regular fungal media (SDY and PDA), which may be the origin of the brownish color when the *MrPks1* was knocked out from the genome.

The last missing puzzle of the DHN melanin pathway in *M. robertsii* was the oxidative enzyme that polymerizes DHN. According to the synteni analysis, MLac1 in MrPKS1 BGC can be one candidate. However, co-expressing the MrPKS2 BGC with *MLac1* did not produce new product peak after 36-h or 48-h fermentation (Fig. 4A). When expressing *MrPks2*, the yeast fermentation was dark brown, in accordance with the color of flaviolin. Sequential addition of *MrTHNR* and *MrSCD* to the pathway lightened the color to be reddish-brown. The color of the fermentation (Fig. 4B) did not change significantly when the MLac1 functioned. To further verify this result, DHN compound was fed to the yeast expressing *Mlac1*, which displayed similar product profile with the yeast control expressing empty plasmid (Fig. S6). Thus, the DHN product could not be transformed by *MLac1* in yeast.

### 3.6 The expression of the MrPks1 gene clusters

The pigment pathway following MrPKS1 is more complicated than that of MrPks2. According to the synteni analysis and previous study, there are at least two possibilities: 1) the product **1** merges into the DHN pathway through genes in MrPks2 BGC; 2) **1** diverged into a different pathway utilizing MLac1 and MrEthD. Based on the RT-PCR analysis (Fig. S4), the first hypothesis is not likely because that *MrTHNR* and *MrSCD* were not activated on regular incubation conditions. As expected, the combinatorial expression of *MrPks1* with *MrTHNR* and/or *MrSCD* failed to further modify **1** (Fig. S7).

To test the second hypothesis, we co-expressed *MrPks1* with combinations of *MLac1* or *MrEthD* genes both cloned from the cDNA of *M. robertsii* (Fig. 5). MrLac1 did not accept **1-3** as substrate (Fig. 5). Addition of MrEthD eliminated **1**, the major product of MrPKS1, while significantly increased the yield of **2** and **3** (Fig. 5D). No new products were detected. The color of the fermentation did not change significantly when adding MrEthD and/or MLac1 to the pathway. After individually fed to the yeast expressing *MrEthD*, **1** disappeared from the product profile without new product (Fig. S7). This result showed that **1** was the substrate of MrEthD and was transformed into a form that cannot be extracted by organic solvent. According to previous study, MLac1 also contributes to the pigmentation of *Metarhizium* fungi.^29, 46^ It is reasonable to hypothesize that Mlac1 can polymerize the activated form of **1** by MrEthD. However, further addition of MLac1 to MrPKS1 and MrEthD did not produce new product peak, nor changed the color of the yeast fermentation in yeast (Fig. 5).

**Figure 5.**
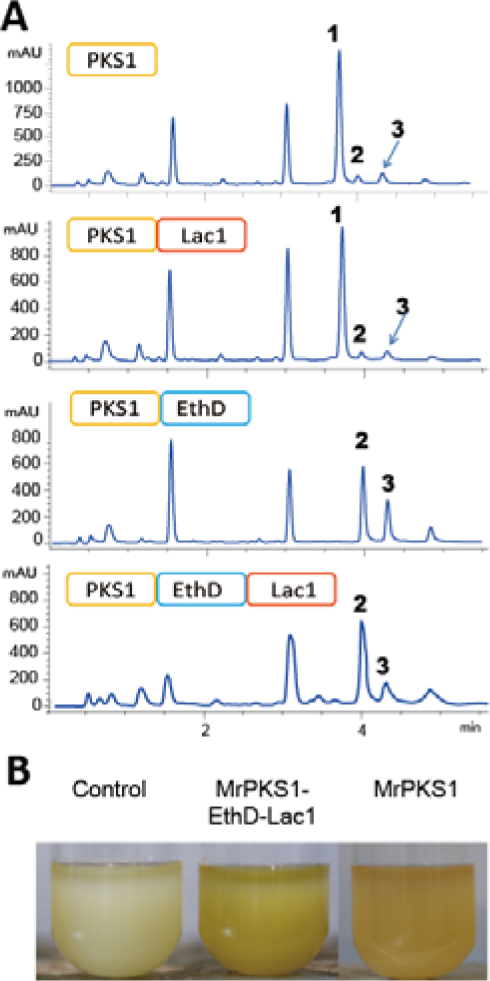
Reconstruction of MrPKS1 pathway in yeast. A. Product profiles (reversed phase HPLC-UV traces recorded at 280 nm) of *S. cerevisiae* BJ5464-NpgA expressing the indicated genes from the MrPKS1 gene cluster. B. The colors of yeast fermentations expressing indicated genes after 48 hr.

### 3.7 Biological function of the MrPKS1 and MrPKS2 products

The intrinsic reason why the pathogenic fungi developed two different melanin biosynthesis pathways is an interesting topic in understanding the fungal adaptation to host. The polymerized DHN melanin has been extensively proved to protect fungal cells from biotic and abiotic stresses, especially important to pathogenic fungi. In this study, the small molecule intermediates, i.e., DHN and T4HN (flaviolin), also can protect the yeast cells against UV-B radiation to different extend (Fig. 6). The yeast producing T4HN (expressing *MrPks2*) showed strongest UV-B radiation resistance than the others under 30-min (0.36 J cm^-2^) dose, while the yeast expressing *MrPks2* along with *MrTHNR* showed weaker protection effect. Similarly, the yeast expressing *MrPks1* or *MrPks1* and *MrEthD* also had stronger resistance to UV-B radiation, showing the ability of the intermediate products of MrPks1 BGC to protect cell against this stress. In contrast, the MrPks1 or MrPks2 pathway products did not show protective affect to yeast cells under heat stress (Fig. S8). This experiment was repeated three times with two different clones each time and similar trends were observed.

**Figure 6.**
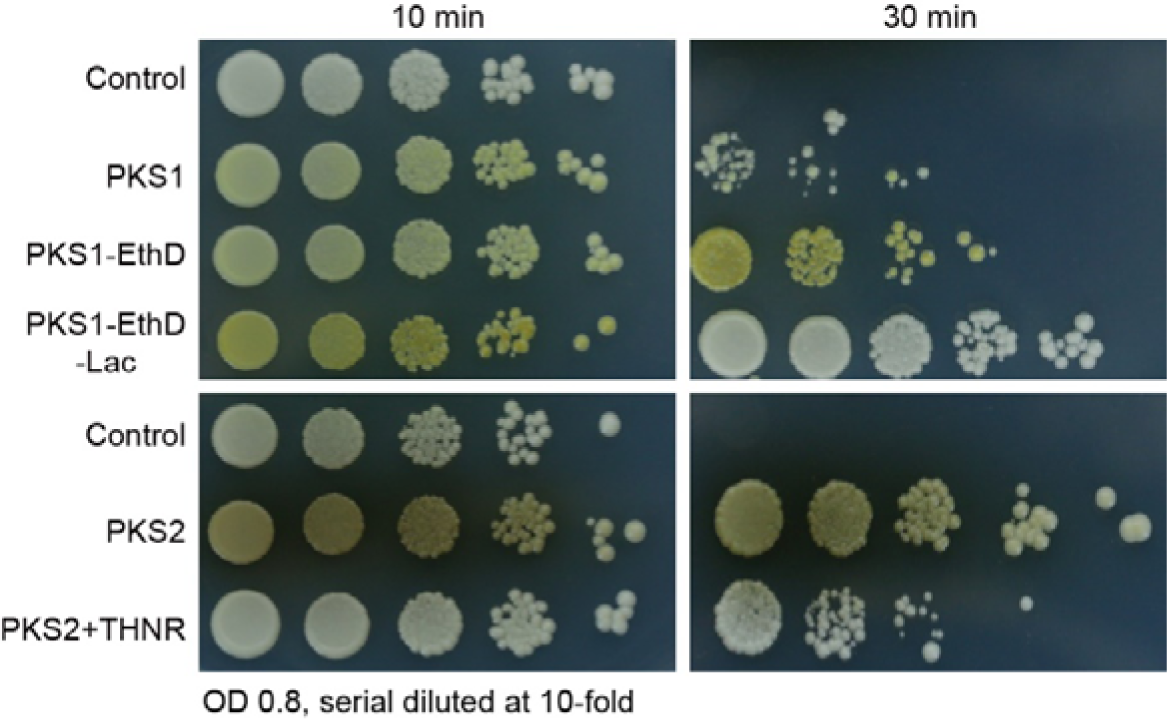
The UV-B stress resistance of yeast expressing the genes in MrPKS1 or MrPKS2 gene clusters. The short names of the heterologously expressed genes were listed left to the photos. The yeast transformants all contains three plasmids with the control expressing three empty plasmids. Stress conditions were marked above the photos.

## 4. Discussion

Many fungi are melanized and the production of the dark pigment can protect the organisms from diverse environmental stresses. For example, melanins confer resistance to UV light by their ability to absorb a broad range of the electromagnetic spectrum and thus prevent photo-induced damage.^3^ In pathogenic fungi, melanins also protect the fungus from oxidative attack by neutrophils, and consequently increase the invasiveness of fungi by the escape of the host immune system.^47^ The wide existence and large quantities as a metabolic product of melanin demonstrate the importance of fungal melanization. DHN melanin is among the most important type of melanin pigment. Fungi have developed different DHN biosynthesis pathways (Fig. 7). Even in the same strain such as *M. robertsii*, at least two divergent pathways have been developed to make sure the melanization is fully functioning during different growth stages. In this study, we confirmed the existence of DHN-biosynthesis pathway in *M. robertsii* by reconstruction of the full pathway in heterologous host, *Saccharomyces cerevisiae* BJ5464-NpgA. Its core biosynthetic enzyme, the polyketide synthase MrPKS2, directly produces T4HN without the assistance of serine hydrolase to remove the acetyl group (Fig. 7). Another polyketide synthase, MrPKS1 is highly similar to MrPKS2 and resulted from duplication event.^28^ Its main product is a new octaketide, which is similar to but different from T4HN with three extra acetyl groups attached to the naphthopyrone ring. This product, **1**, has high reactivity and can be transformed by yeast or *A. nidulans*, or even spontaneously to different forms (**2-4**). It cannot be transformed by the MrTHNR and MrSCD enzymes to produce DHN; instead, it is very likely to be activated by the MrEthD encoded adjacent to the *MrPks1* gene, and formed a new type of pigment. Other naphthopyranone or anthraquinone compounds synthesized by polyketide synthase also were reported to be polymerized directly to form pigment, serving as alternatives to DHN melanin. For example, YWA1 synthesized by wA of *A. nidus* can form green spore pigment;^48^ The anthraquinone compound asparasone synthesized by pks27 of *A. flavus* can form dark brown pigment, but the specific pathway is still unknown;^49^ similar compounds synthesized by *Paecilomyces vannamei* can also form dimeric viriditoxin through the action of oxidase.^50^ The plethora of these pigments again emphasizes the importance of fungus melanization, and may help the producing strains to adapt to different living environments. For example, DHN melanin has been reported as virulence factors to trigger the host immune response;^11, 47^ thus alternatives may be developed to escape the host immune system. The outstanding protective effects of DHN, its intermediates and the naphthopyrone counterparts against UV-B radiation were demonstrated in this study. The intrinsic physiological reason of this phenomenon would be an interesting topic to discuss further.

## Acknowledgement

This work was supported by the National Key Research and Development Program of China (2023YFF1000300 and 2023YFF1000302 to L.Z.), the National Natural Science Foundation of China (32070064 and 32270069 to L.Z.), fellowship of China National Postdoctoral Program for Innovative Talents (BX20220348 to L.X.), and the Central Public-interest Scientific Institution Basal Research Fund (to Y.X. and L.Z.). We sincerely thank Prof. Chengshu Wang from Center for Excellence in Molecular Plant Sciences, Shanghai Institute of Plant Physiology and Ecology, CAS, for providing the *Metarhizium robertsii* strain and cDNA. We also thank the Beijing Center for Physical and Chemical Analysis for providing access to their NMR device. All authors declare no competing financial interests.

