## Supplementary files for "The divergence of DHN-derived melanin pathways in *Metarhizium robertsii*"

### Structure elucidation

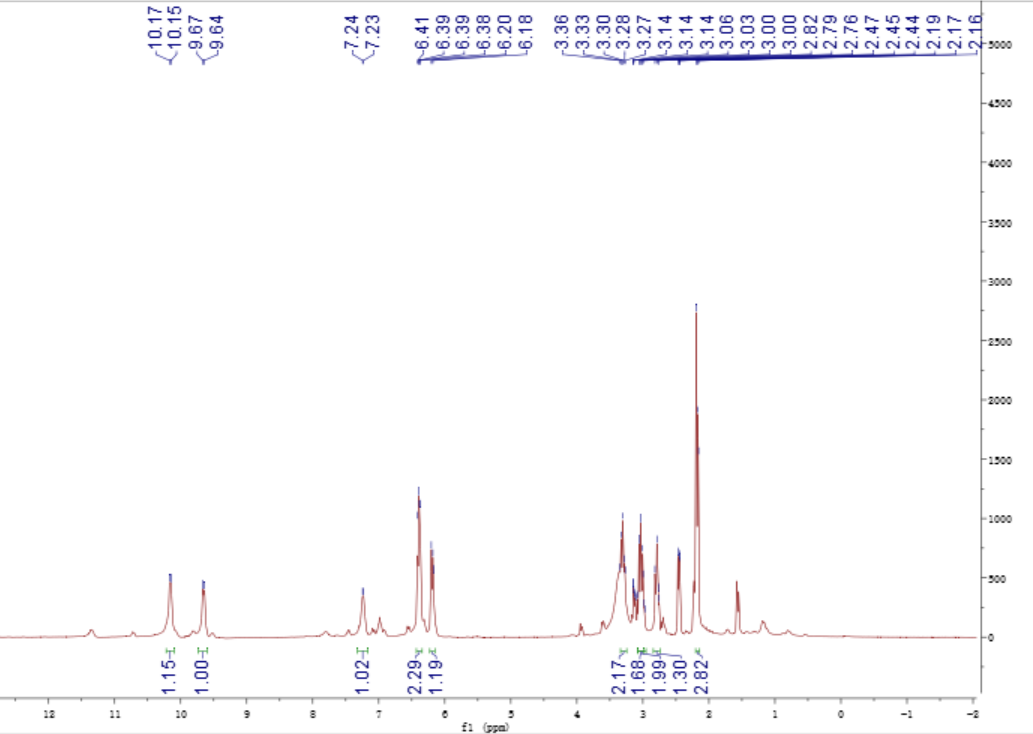

A. ^1^H- NMR (600 MHz) spectrum of **1** in DMSO-*d*_6_

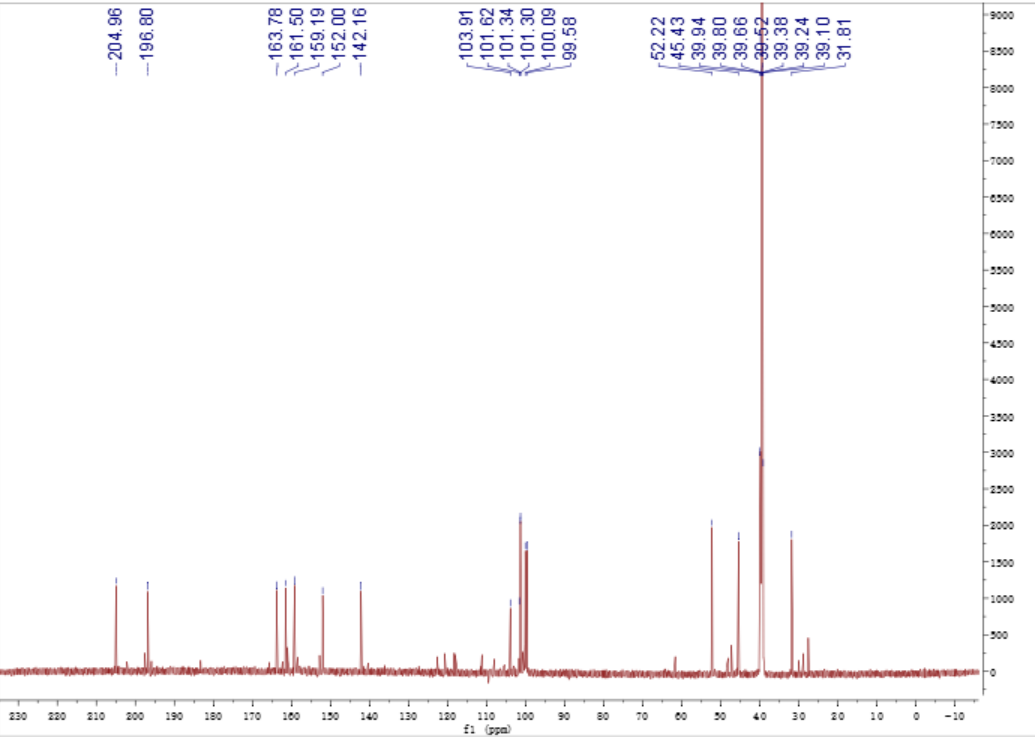

B. ^13^C- NMR (150 MHz) spectrum of **1** in DMSO-*d*_6_

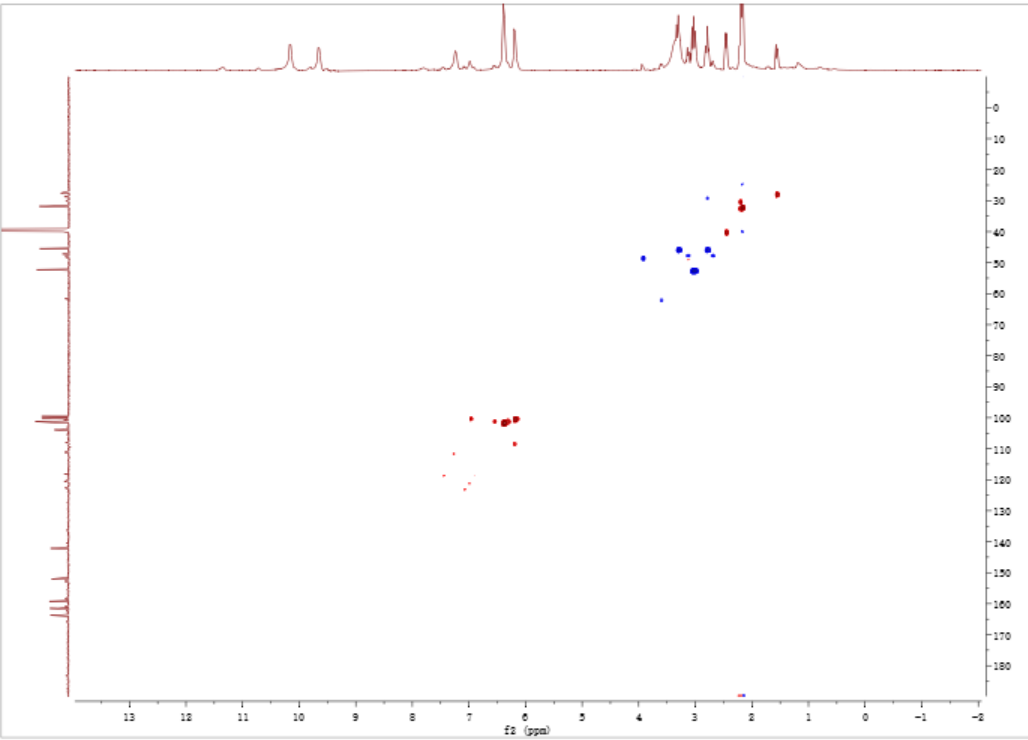

C. HSQC spectrum of **1** in DMSO-*d*_6_

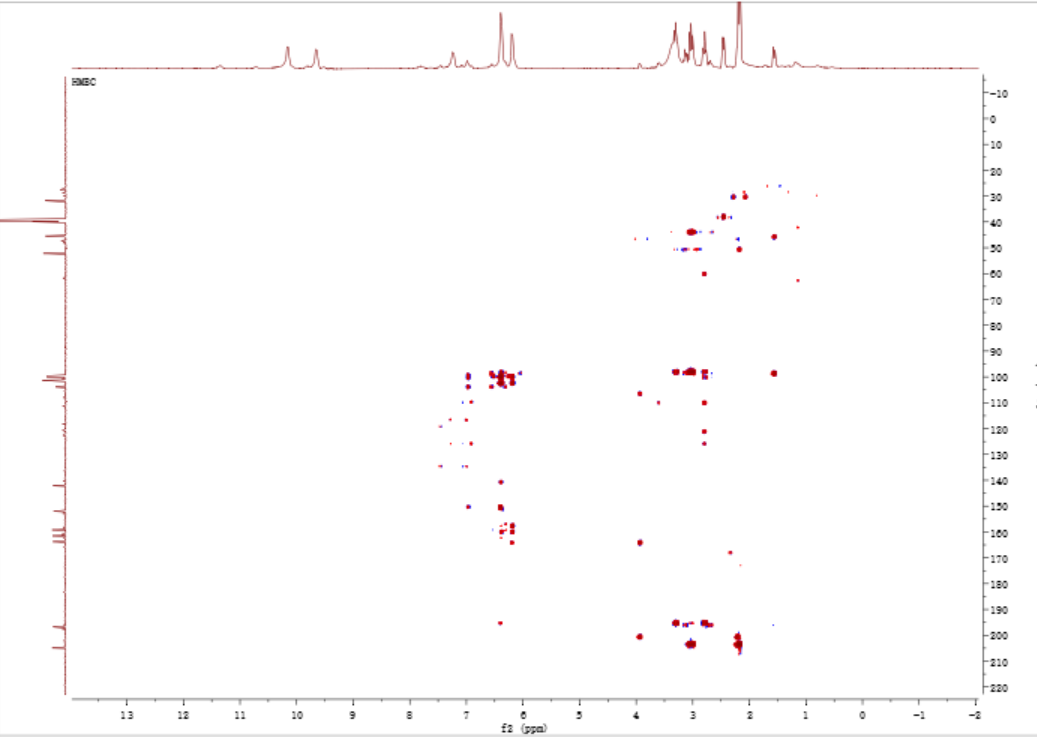

D. HMBC spectrum of **1** in DMSO-*d*_6_

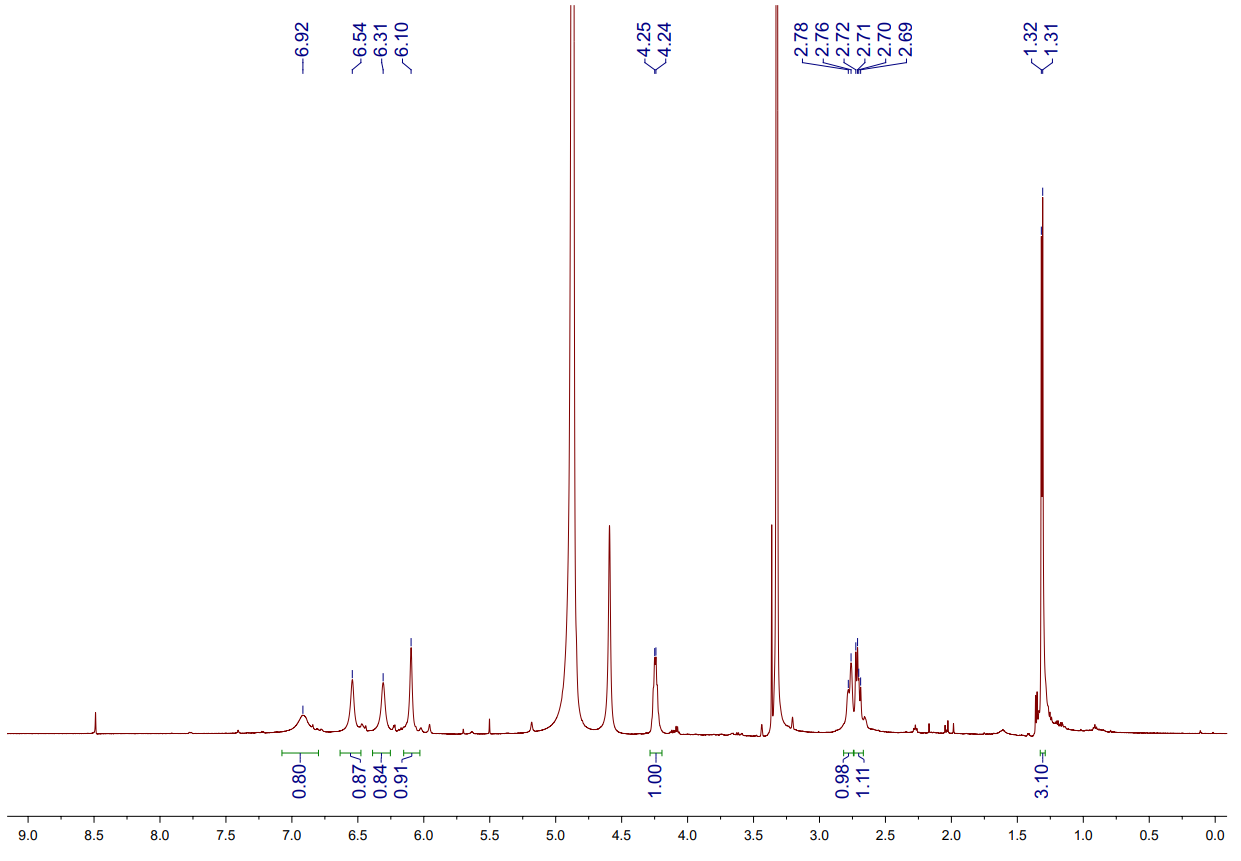

E. ^1^H NMR spectrum of **2** in methanol-*d*_4_

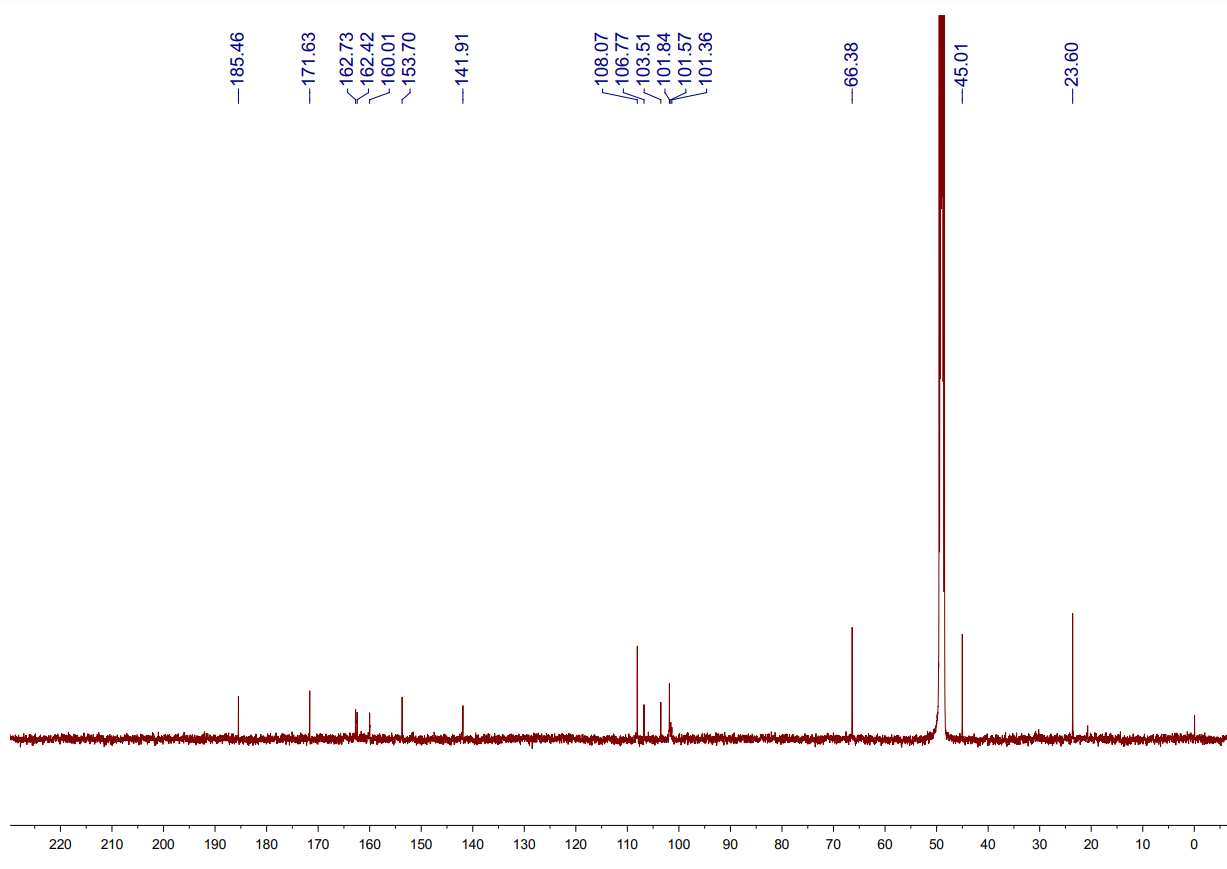

F. ^13^C NMR spectrum of **2** in methanol-*d*_4_

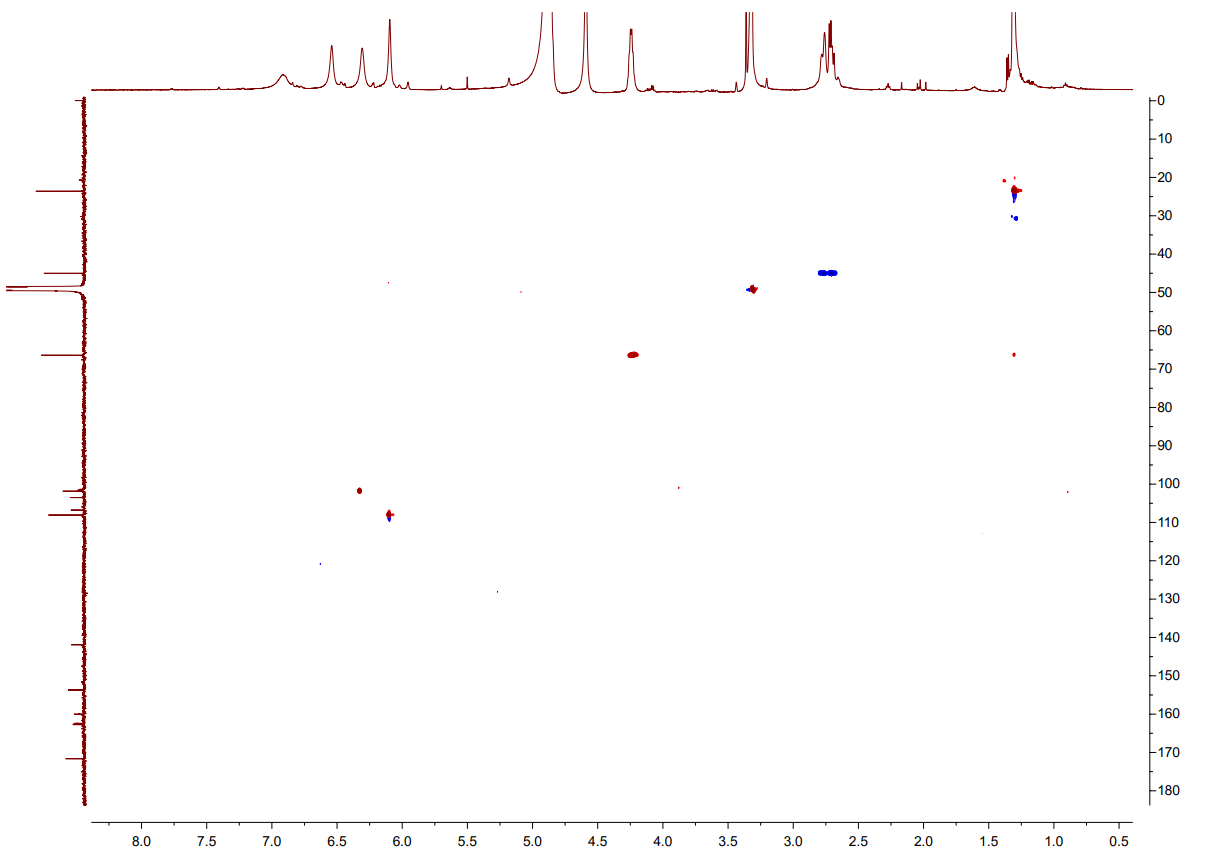

G. HSQC spectrum of **2** in methanol-*d*_4_

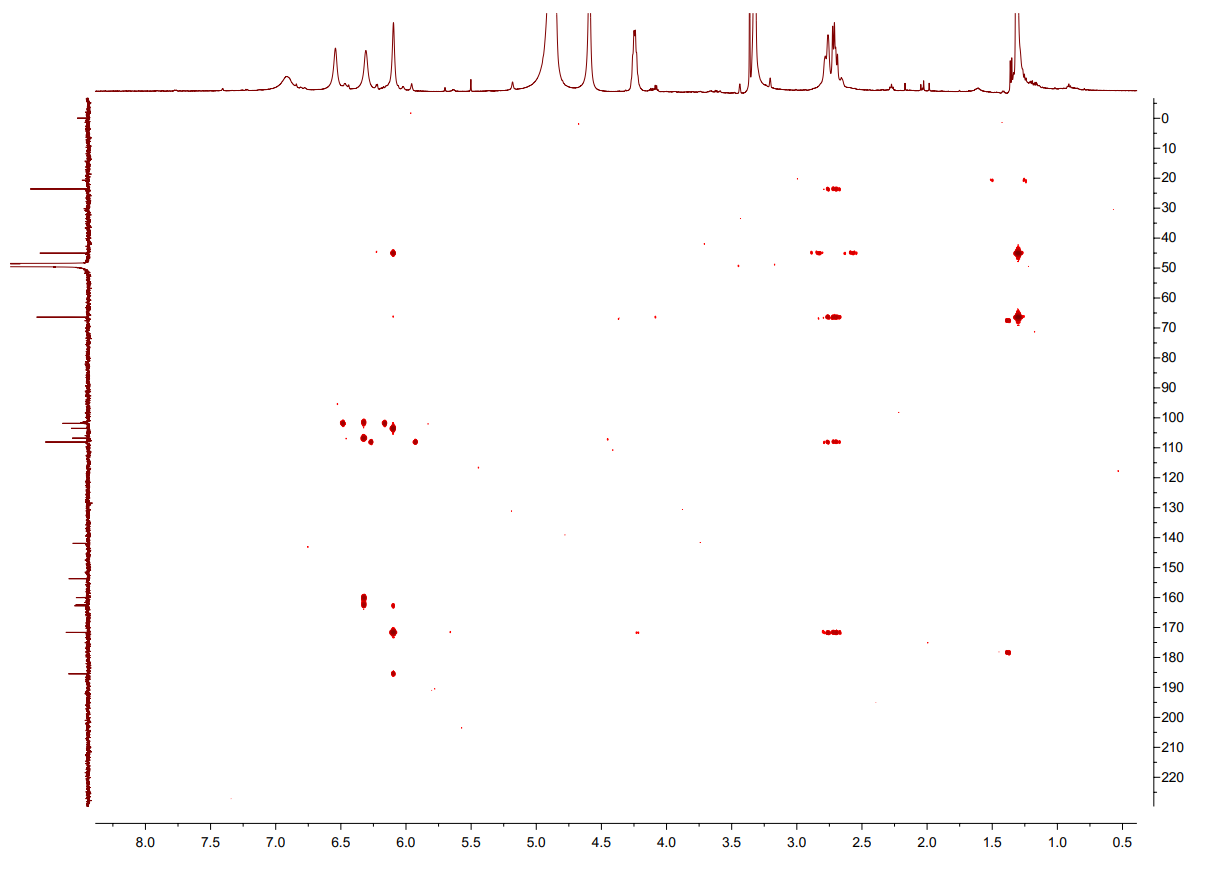

H. HMBC spectrum of **2** in methanol-*d*_4_

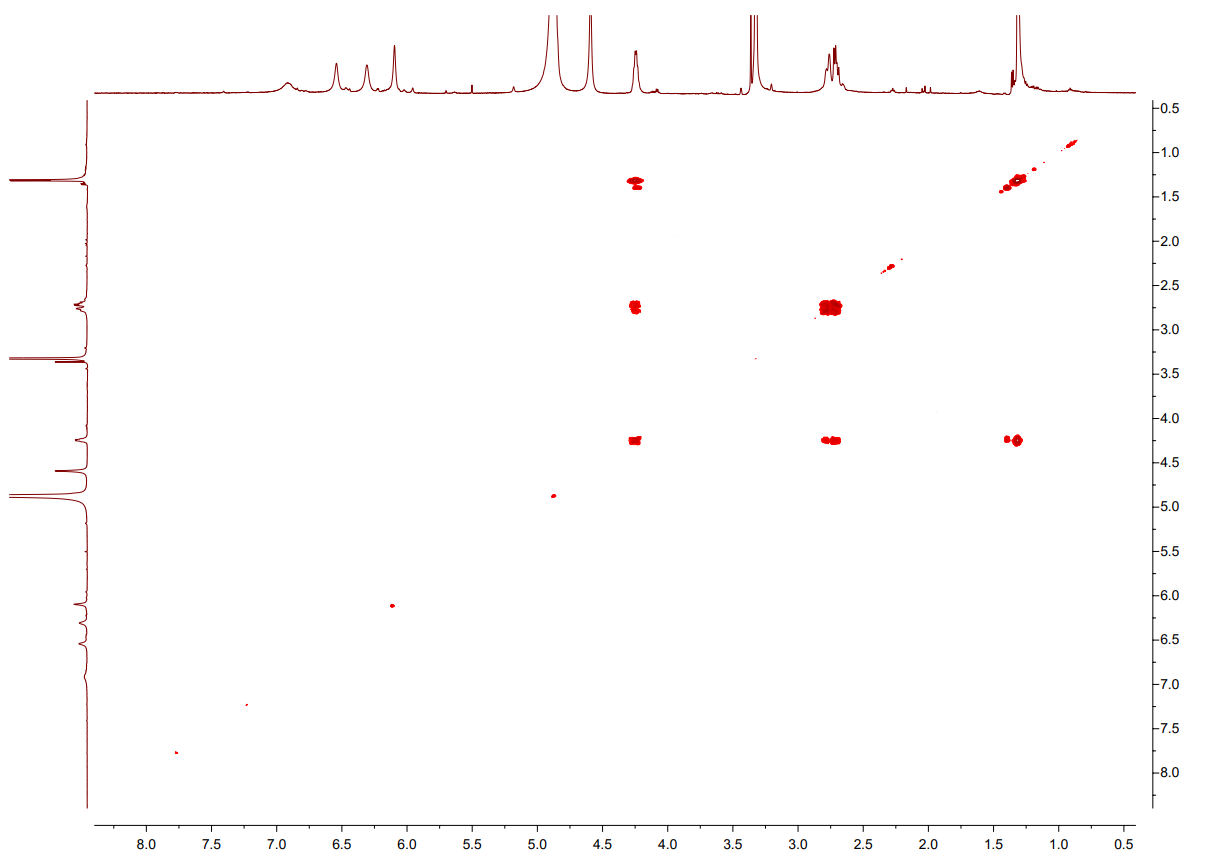

I. ^1^H-^1^H COSY spectrum of **2** in methanol-*d*_4_

Figure S1. The NMR spectra of **1** (A-D) and **2** (E-I)**.**

#### Table S1. ^1^H- (600 MHz) and ^13^C-NMR (150 MHz) Data of **1**, **2** and **3** in MeOD-*d_6_* or DMSO-*d_6_*

A. ^1^H- (600 MHz) and ^13^C-NMR (150 MHz) Data of **1** in DMSO-*d*_6_

| **No.** | **1** | |
| --- | --- | --- |
|  | ***δ*_C_** | ***δ*_H_ (multi, *J* in Hz)** |
| 2 | 100.1 |  |
| 3*α* | 45.4 | 3.29, m |
| 3*β* |  | 2.78, m |
| 4 | 196.8 |  |
| 4a | 101.6 |  |
| 5 | 163.8 |  |
| 5a | 103.9 |  |
| 6 | 151.2 |  |
| 7 | 100.1 | 6.19, s |
| 8 | 161.5 |  |
| 9 | 101.3 | 6.39, s |
| 9a | 142.2 |  |
| 10 | 101.3 | 6.39, s |
| 10a | 152.0 |  |
| 11 | 52.7 | 3.02, m |
| 12 | 205.0 |  |
| 13 | 31.8 | 2.17, s |

B. ^1^H- (600 MHz) and ^13^C-NMR (150 MHz) Data of **2** in methanol-*d*_4_

| **No.** | **2** | |
| --- | --- | --- |
|  | ***δ*_C_** | ***δ*_H_ (multi, *J* in Hz)** |
| 2 | 171.6 |  |
| 3 | 108.1 | 6.10, s |
| 4 | 185.5 |  |
| 4a | 103.5 |  |
| 5 | 162.7 |  |
| 5a | 101.6 |  |
| 6 | 160.0 |  |
| 7 | 101.8 | 6.31, s |
| 8 | 162.4 |  |
| 9 | 106.8 | 6.54, s |
| 9a | 141.9 |  |
| 10 | 101.4 | 6.92, s |
| 10a | 153.7 |  |
| 11*α* | 45.0 | 2.71, dd (13.7, 8.2) |
| 11*β* |  | 2.77, br d (13.6) |
| 12 | 66.4 | 4.24, m |
| 13 | 23.6 | 1.31, d (6.1) |

#### Table S2. The *m*/*z* values of parent ions and major daughter ions of compounds **1**-**3** in positive ion mode.

| **Compound** | **Parent ion *m*/*z* (amu)** | **Calculated *m*/*z* (amu)** | **Mass error (ppm)** | **Major daughter ion *m*/*z* (amu)** |
| --- | --- | --- | --- | --- |
| **1** | 319.0784 | 319.0817 | 10.30 | 219.0187, 259.0510, 301.0728 |
| **2** | 303.0871 | 303.0869 | 0.66 | 259.0602, 285.0753 |
| **3** | 301.0727 | 301.0712 | 4.98 | 259.0616 |

#### NMR analysis of compound **2**

The ^1^H NMR data for compound **2** (Table S1-2) revealed the presence of four unsaturated hydrogen proton signals [*δ*_H_ 6.10 (1H, s), 6.31 (1H, s), 6.54 (1H, s), 6.92 (1H, s)] and one singlet methyl signal [*δ*_H_ 1.31 (1H, d, *J* = 6.1 Hz, H-13)], one methylene signal [*δ*_H_ 2.76 (1H, br d, *J* = 13.6 Hz, H-11*β*), 2.71 (1H, dd, *J* = 13.7, 8.2 Hz, H-11*α*)], one oxymethine [*δ*_H_ 4.24 (1H, m, H-12)]. The ^13^C NMR spectrum (SI Table S1-2), with the aid of HSQC, showed 16 resonances ascribed to one methyl, one methylene, five methines, and nine quaternary carbons. Analysis of 1D and 2D NMR data (Fig. S1 and Table S1) indicated that compounds **1** and **2** share the same aromatic ring skeleton.

### The reactivity of MrPKS1 products

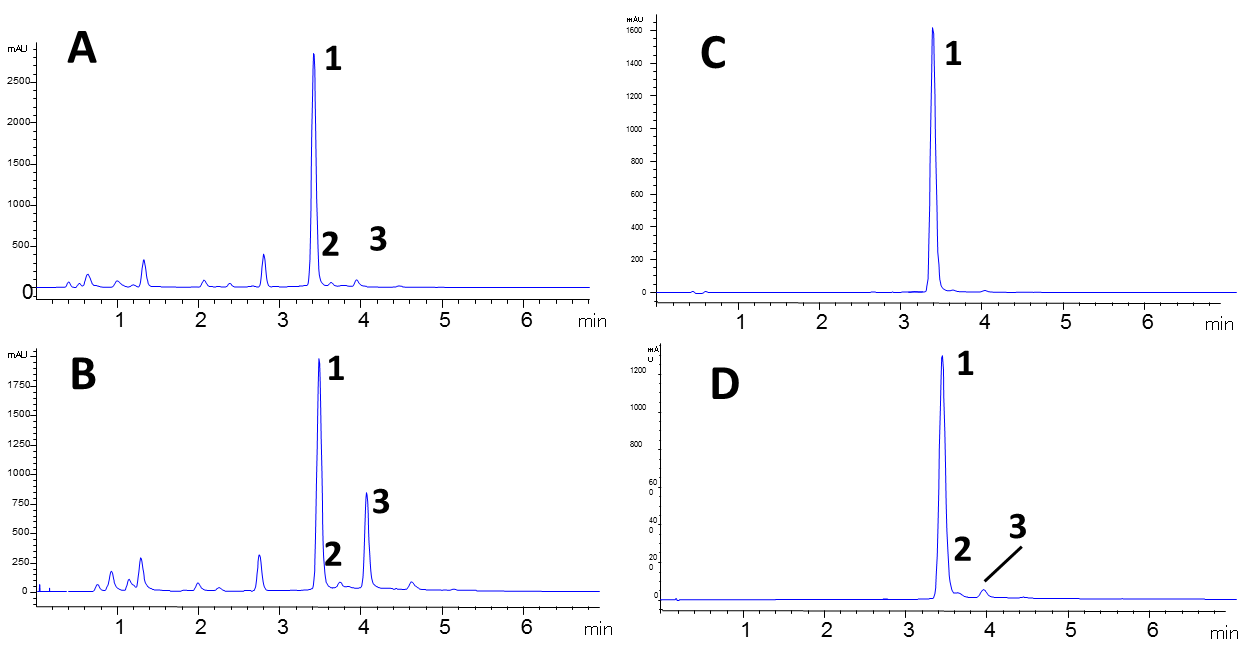

Figure S2. The spontaneous transformation of 1 to 2 and 3: time course of. A. MrPKS1 product profiles (reversed phase HPLC-UV traces recorded at 280 nm) in crude extract after 36 hr fermentation. B. The same extract of A stored in room temperature after one week. C. The product profiles of purified **1** recorded immediately and D. after one month.

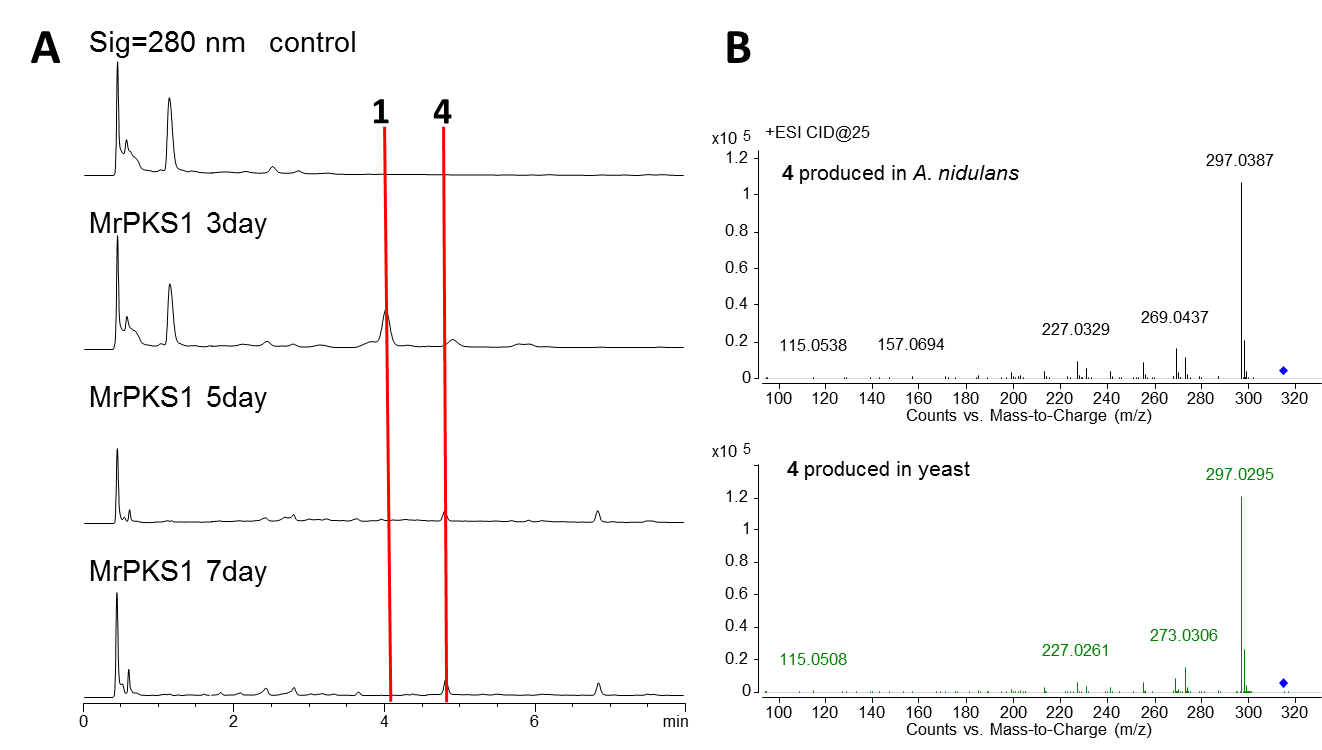

Figure S3. A. The time course of MrPKS1 product profiles (reversed phase HPLC-UV traces recorded at 280 nm) expressed in *A. nidulans* after 3, 5 or 7 d fermentation. B. The comparison of MS/MS spectra of 4 produced in *A. nidulans* and yeast expressing *MrPks1* gene.

Table S3. Genbank accession number for the T4HN PKS and their fungal species.

| Fungal species | Accession of PKS genes |
| --- | --- |
| *Colletotrichum lagenarium* (*CIPks1*) | TDZ26149.1 |
| *Verticillium dahliae* | XP_009649862.1 |
| *Sporothrix schenckii* | ERS95912.1 |
| *Pyricularia grisea* | TLD09340.1 |
| *Botrytis cinerea* (*bcpks12*) | XP_024547374.1 |
| *Hortaea werneckii* | RMY92220.1 |
| *Paracoccidioides brasiliensis* | ODH34604.1 |
| *Aspergillus fumigatus* (*Alb1*) | XP_756095.1 |
| *Aspergillus nidulans* (*wA*) | CBF74114.1 |
| *Metarhizium acridum* (*McPks2*) | XP_007815650.1 |
| *Metarhizium anisopliae* (*MnPks2*) | KID71243.1 |
| *Metarhizium roberstii* (*MrPks2*) | EFZ02010.2 |
| *Fusarium graminearum* | XP_011318224.1 |
| *Metarhizium acridum* (*McPks1*) | XP_007811725.1 |
| *Metarhizium anisopliae* (*MnPks1*) | KID59669.1 |
| *Metarhizium roberstii* (*MrPks1*) | EFY96684.2 |
| *Botrytis cinerea* (*bcpks13*) | XP_001547095.2 |
| *Wangiella dermatitidis* (*WdPks1*) | EHY55015.1 |
| *Pestalotiopsis fici* ( *PfmaE* ) | XP_007833873.1 |

### The biosynthesis of DHN by the *MrPks2* gene cluster under strict regulation

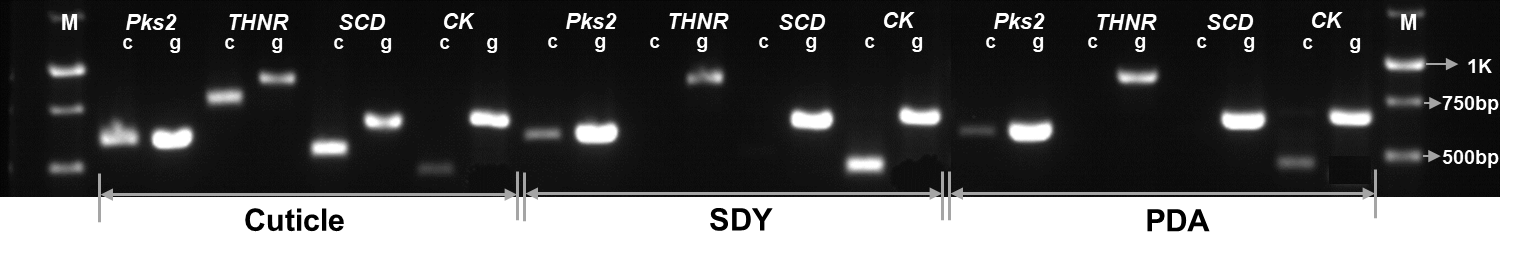

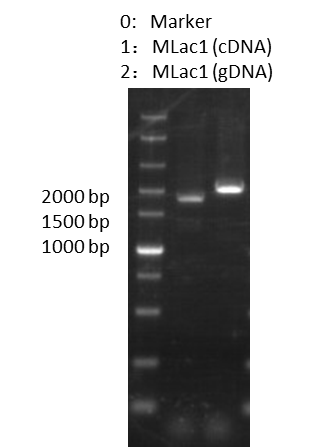

Figure S4. Expression of genes related to DHN biosynthetic pathway on insect cuticle, SDY or PDA media.

c: cDNA, g: genome DNA, THNR: THN reductases, SCD: scytalone dehydratases, CK: tubulin protein.

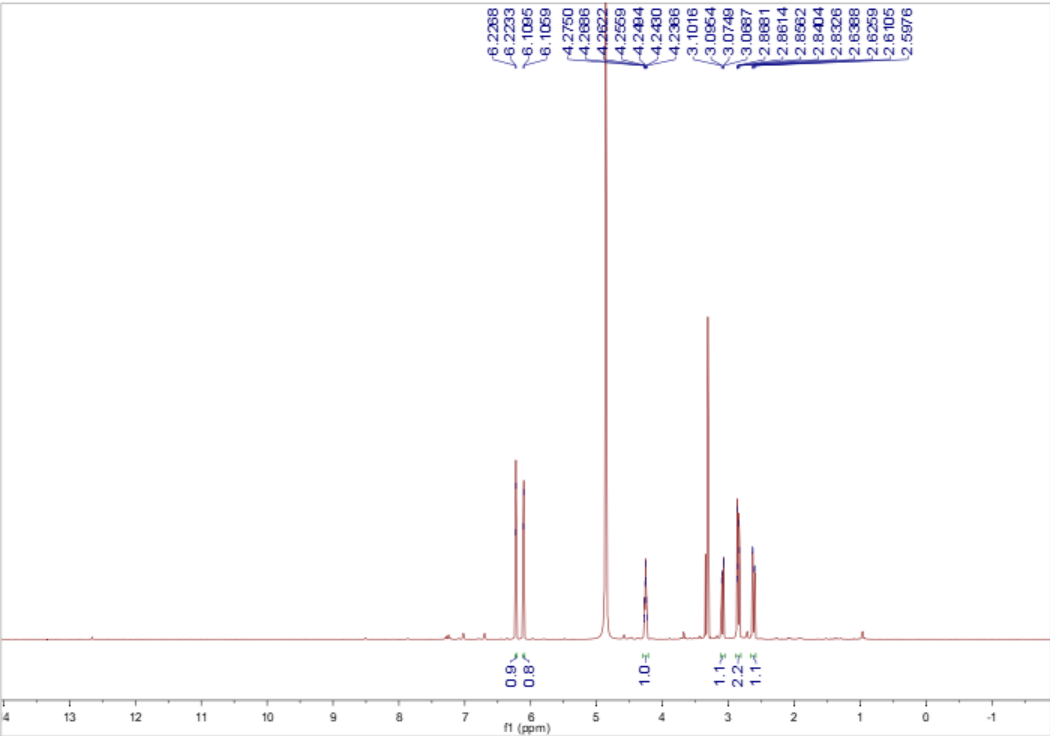

A. ^1^H- NMR (600 MHz) spectrum of scytalone in MeOD.

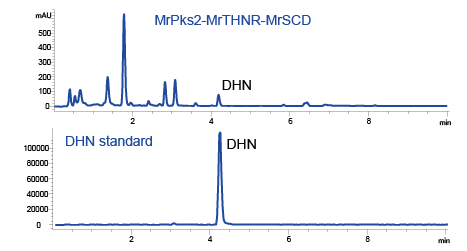

B. the HPLC graphs of DHN standard and the extract of yeast expressing *MrPks2*, *MrTHNR* and *MrSCD*.

Figure S5. A. The NMR spectra of scytalone; B. the HPLC graphs of DHN standard and the extract of yeast expressing *MrPks2*, *MrTHNR* and *MrSCD*.

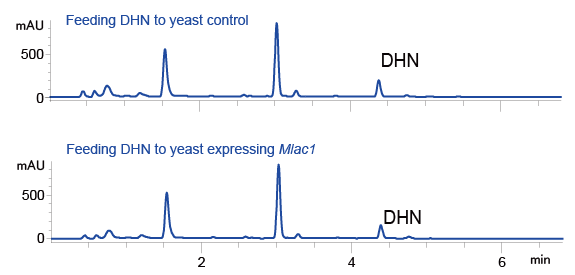

Figure S6. The HPLC graphs of DHN feeding to yeast expressing MLac1or empty plasmid.

Table S4. ^1^H- (600 MHz) and ^13^C-NMR (150 MHz) Data of scytalone in MeOD

A. ^1^H- (600 MHz) and ^13^C-NMR (150 MHz) Data of **Scytalone** in MeOD

| **No.** | **Scytalone** | |
| --- | --- | --- |
|  | ***δ*_C_** | ***δ*_H_ (multi, *J* in Hz)** |
| 1 | 202.4 |  |
| 2*α* | 47.4 | 2.82, dd (17.0, 4.4) |
| 2*β* |  | 2.60, dd (17.0, 7.7) |
| 3 | 66.9 |  |
| 4*α* | 39.1 | 3.07, dd (15.9, 3.4) |
| 4*β* |  | 2.82, dd (15.9, 2.7) |
| 4a | 146.0 |  |
| 5 | 101.6 | 6.09, d (1.7) |
| 6 | 166.5 |  |
| 7 | 109.4 | 6.21, d (1.7) |
| 8 | 166.6 |  |
| 8a | 111.7 |  |

### The expression of the MrPKS1 gene clusters

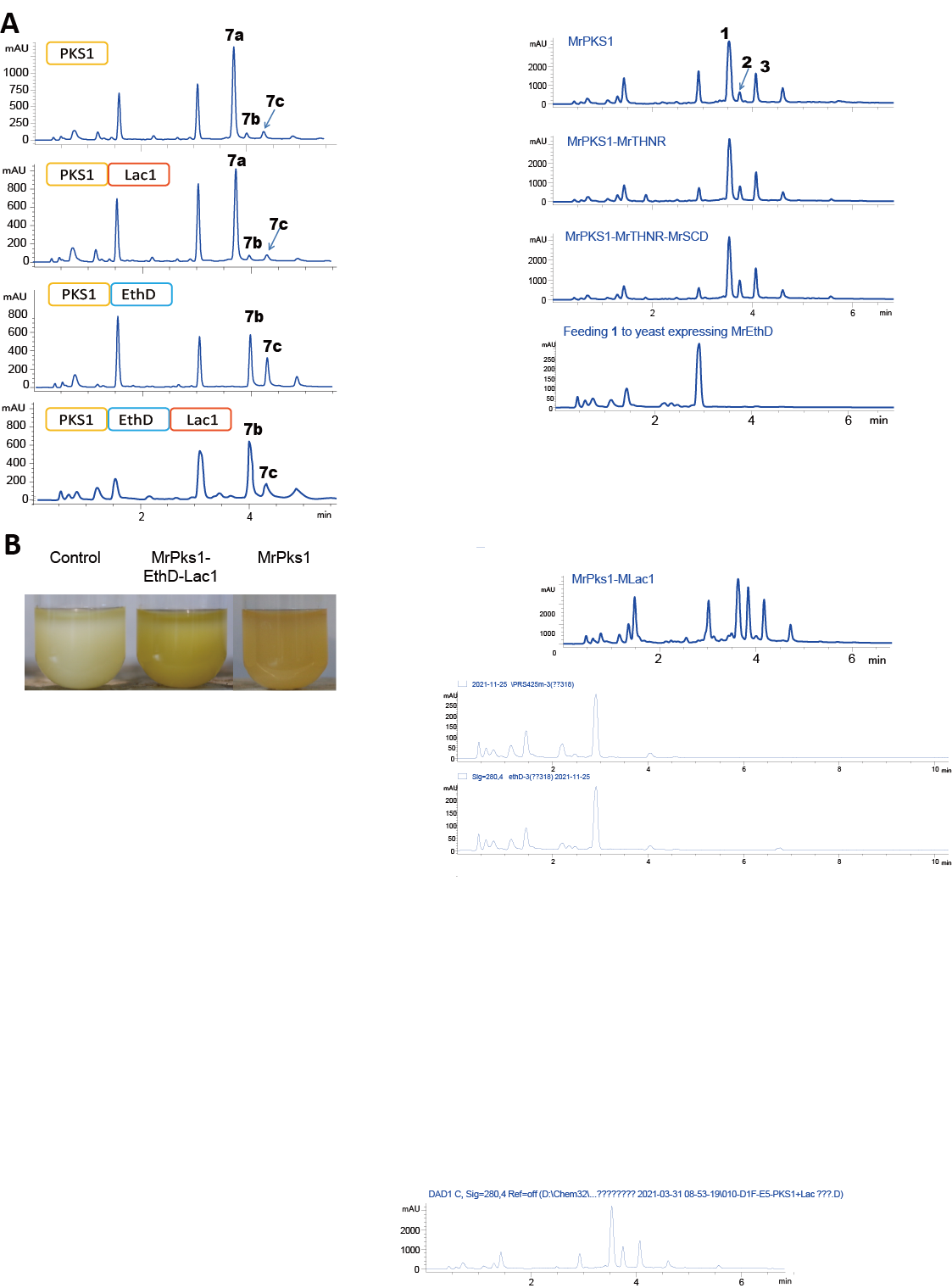

Figure S7. Product profiles (reversed phase HPLC-UV traces recorded at 280 nm) of *S. cerevisiae* BJ5464-NpgA expressing the indicated genes from *M. robertsii*.

### Biological function of the MrPKS1 and MrPKS2 products

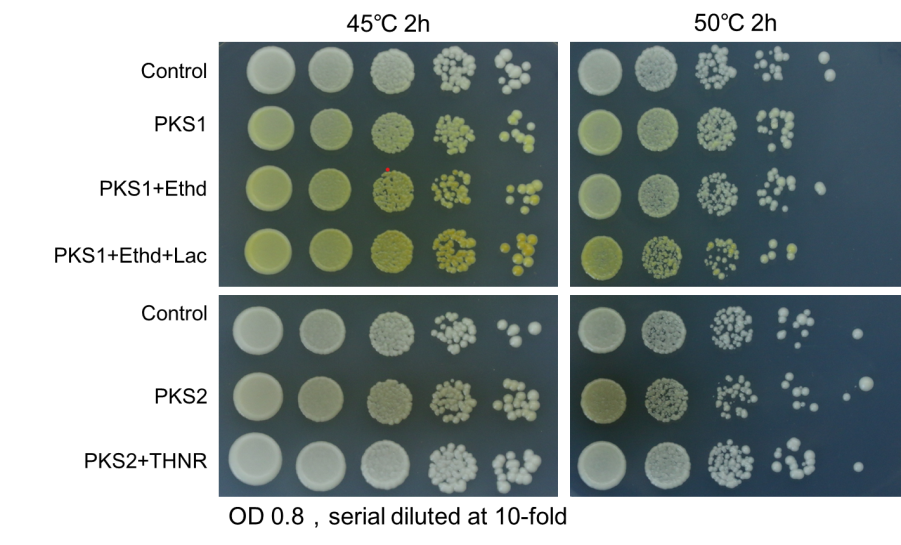

Figure S8. The heat stress resistance of yeast expressing the genes in MrPKS1 or MrPKS2 gene clusters. The short names of the heterologously expressed genes were listed left to the photos. The yeast transformants all contains three plasmids with the control expressing three empty plasmids. Stress conditions were marked above the photos. This experiment was repeated three times with two different clones each time.

### Primer table

Table S5 The primers used in this study

| Primer | | | Sequence (5′-3′) | Length (bp) |
| --- | --- | --- | --- | --- |
| Laccase-1F | | GATGACGACAAGCTTCATATGAGCCGCTTTGCGCGTC | | 37 |
| Laccase-1R | ATGGTGATGTCCGTTTAAACCTAACGCAGCTGATATTCTTCAGG | | | 44 |
| THNR-1F | GATGACGACAAGCTTCATATGGCATCCAGCGAGGAA | | | 36 |
| THNR-3R | ATGGTGATGTCCGTTTAAACTTAGGTGACGGAGCCTCCAGTAAGGAGGATAACTTGGC | | | 58 |
| SCD-1F | GATGACGACAAGCTTCATATGAACCAATCTTTAACCCC | | | 38 |
| SCD-1R | ATGGTGATGTCCGTTTAAACCTACGAATGTTTCTCCGACTG | | | 41 |
| EthD-F | GATGACGACAAGCTTCATATGATGGCTATTTCTGATTC | | | 38 |
| EthD-R | TGGTGATGGTGATGTCCGTTTAAACTTATTTTGGAACAAC | | | 40 |
| PKS1-2-F | GACCATGATTACGCCA | | | 16 |
| PKS1-TE-R | TTAACCAAACFTACTCCGAATA | | | 22 |
| PKS2-2-F | CGCAACGCAATTAATG | | | 16 |
| PKS2-TE-R | CTAGCTGGGACCAAGCAAAAATTC | | | 24 |
